# Context-sensitive processing in a model neocortical pyramidal cell with two sites of input integration

**DOI:** 10.1101/2024.01.16.575982

**Authors:** Bruce P. Graham, Jim W. Kay, William A. Phillips

**Affiliations:** Computing Science and Mathematics, Faculty of Natural Sciences, University of Stirling, Stirling, U.K.; School of Mathematics and Statistics, University of Glasgow, Glasgow, U.K.; Psychology, Faculty of Natural Sciences, University of Stirling, Stirling, U.K.

**Keywords:** pyramidal cells, burst firing, contextual processing, partial information decomposition, transfer functions

## Abstract

Neocortical layer 5 thick-tufted pyramidal cells are prone to exhibiting burst firing on receipt of coincident basal and apical dendritic inputs. These inputs carry different information, with basal inputs coming from feedforward sensory pathways and apical inputs coming from diverse sources that provide context in the cortical hierarchy. We explore the information processing possibilities of this burst firing using computer simulations of a noisy compartmental cell model. Simulated data on stochastic burst firing due to brief, simultaneously injected basal and apical currents allows estimation of burst firing probability for different stimulus current amplitudes. Information-theory-based partial information decomposition (PID) is used to quantify the contributions of the apical and basal input streams to the information in the cell output bursting probability. Four different operating regimes are apparent, depending on the relative strengths of the input streams, with output burst probability carrying more or less information that is uniquely contributed by either the basal or apical input, or shared and synergistic information due to the combined streams. We derive and fit transfer functions for these different regimes that describe burst probability over the different ranges of basal and apical input amplitudes. The operating regimes can be classified into distinct modes of information processing, depending on the contribution of apical input to output bursting: *apical cooperation*, in which both basal and apical inputs are required to generate a burst; *apical amplification*, in which basal input alone can generate a burst but the burst probability is modulated by apical input; *apical drive*, in which apical input alone can produce a burst; and *apical integration*, in which strong apical or basal inputs alone, as well as their combination, can generate bursting. In particular, PID and the transfer function clarify that the apical amplification mode has the features required for contextually-modulated information processing.

## 1 Introduction

One of the key challenges in understanding cortical information processing is in building a comprehensive picture of pyramidal cells as two-point processors receiving dual information-rich input streams that must be combined to produce an output that is part of a meaningful and coherent pattern of activity across the system or sub-system of which those cells are a part. Pyramidal cells are distinctly not single-point processors, as commonly used in artificial neural networks. Neocortical pyramidal cells have essentially two sites of synaptic integration, targetting the basal dendrites and apical tuft dendrites, respectively (Larkum, 2013; Phillips, 2017). The basal inputs are in the perisomatic region and so arrive close to the final site of cell synaptic integration and action potential initiation. These inputs are from feedforward sources, conveying sensory information via specific regions of the thalamus and cortex and provide the primary drive to the receiving pyramidal cell that determines its receptive field. On the other hand, inputs to the apical tuft come from diverse feedback sources in higher cortex, long-range lateral cortex and thalamus. These inputs are hypothesised to provide contextual information that can modulate the cell’s response to its primary drive (Phillips, 2017). This has been termed *apical amplification* (Phillips, 2017; Phillips et al., 2018; Marvan et al., 2021; Marvan and Phillips, 2024; Phillips, 2023). Contextually-modulated information processing is widespread in the neocortex and underpins many cognitive functions during conscious processing (Phillips et al., 2015; Aru et al., 2020b; Marvan et al., 2021; Marvan and Phillips, 2024; Phillips, 2023).

A candidate cellular mechanism for apical amplification is present in thick-tufted layer 5 pyramidal cells. In these cells the apical tuft is particularly electrically isolated from the soma, but it contains its own initiation zone for broad calcium spikes. Such spikes are capable of generating a bursting output of sodium spikes from the cell. Apical calcium spikes are often initiated by a combination of apical synaptic input with a back-propagating sodium spike initiated somatically by basal inputs. This has been termed backpropagation-activated calcium (BAC) spike firing (Larkum, 2013). At the cell activity level, there is direct experimental evidence from the visual system and the barrel cortex that PC receptive field responses can be altered in task-dependent ways, indicating contextually-modulated processing via apical amplification (Gilbert and Li, 2013; Takahashi et al., 2016; Dadarlat and Stryker, 2017; Pakan et al., 2018).

Other behaviourally-relevant information processing modes have been identified. There is evidence for *apical drive*, in which apical inputs alone can generate spiking output in the pyramidal cell (Aru et al., 2020a; Phillips, 2023). Apical drive is hypothesised to be fundamental to dreaming (Aru et al., 2020a), but is also apparent in the awake animal. Visual activity has been recorded due to locomotion in the dark, when visual sensory information is low (Keller et al., 2012). In the barrel cortex, increased excitability in PC dendrites leads to an increased detection of apparent whisker stimulation in the absence of actual physical stimulation (Takahashi et al., 2016). Another mode is *apical isolation* in which the apical inputs have no effect on cell output, which is purely driven by basal (perisomatic) inputs (Aru et al., 2020a; Phillips, 2023). Apical isolation might be in effect during dreamless sleep (Aru et al., 2020a). General anesthesia has been shown to decouple pyramidal neurons by effectively cutting off the affects of apical input on cell output (Phillips et al., 2018; Suzuki and Larkum, 2020).

Apical inputs are subject to non-linear processing in the dendrites that can lead to dendritic spiking through activation of voltage-gated calcium and ligand-gated NMDA ion channels (Larkum et al., 2022). These dendritic spikes may propagate sufficiently to the soma to increase the cell’s basal-driven firing rate and to create short, high frequency bursts of spikes. Bursting has long been hypothesised to be a powerful information coding signal in the brain (Lisman, 1997; Krahe and Gabbiani, 2004; Zeldenrust et al., 2018). Bursts can very effectively be read out downstream through the filtering of chemical synapses via stochastic transmitter release and short-term plasticity (Lisman, 1997; Naud and Sprekeler, 2018; Zeldenrust et al., 2018). They usually exist within a stream of single spikes and theoretical work has established that bursts and single spikes can coexist and carry different information that can be read out by downstream neurons (Oswald et al., 2004; Naud and Sprekeler, 2018; Williams et al., 2021; Zeldenrust et al., 2018; Friedenberger et al., 2023; Friedenberger and Naud, 2023).

Exactly how apical and basal inputs are integrated and combined to generate output firing in pyramidal cells is a subject of considerable interest, both to the neuroscience and artificial intelligence (AI) communities. The segregation of inputs in dendrites, followed by their integration, has significant information processing implications (Makarov et al., 2023; Pagkalos et al., 2024). The segregation of basal and apical inputs can be utilised to try to solve the credit assignment problem for synaptic weight updates in deep learning algorithms (Guerguiev et al., 2017; Sacramento et al., 2018). Burst-dependent synaptic plasticity in segregated neurons can contribute to credit assignment for deep learning in hierarchical neural networks (Payeur et al., 2021) and has been used for target-based (Capone et al., 2023) and context-association learning (Baronig and Legenstein, 2024) in such networks.

To help shed light on exactly how basal and apical input streams may combine to modulate and refine cellular receptive field responses, here we characterise the integration of apical and basal (perisomatic) inputs in a computational model of a thick-tufted layer 5 neocortical pyramidal cell (Bahl et al., 2012), and interpret what this means for information processing. We focus on bursting output as the distinctive signature of the combined effects of coincident basal and apical input. Computer simulations are used to collect data on the bursting output of the model cell for varying strengths of coincident basal and apical inputs. We use information theoretical analysis and transfer function fitting to characterise the information transmitted by bursts about the dual input streams. Partial information decomposition (PID) (Williams and Beer, 2010) is able to quantify the contributions of the apical and basal input streams to the information in the cell output. Depending on the relative strengths of the input streams, output burst probability carries more or less information that is uniquely contributed by either the basal or apical input, or shared and synergistic information due to the combined streams. We derive transfer functions for burst probability that make explicit the functional relationships between the two input streams and bursting output.

By analysing information transmission over different ranges for the strengths of basal and apical inputs we find distinct modes of operation. In addition to apical amplification and apical drive, we identify two further modes: *apical cooperation* in which both basal and apical input is required to generate bursting; and *apical integration* in which strong apical or basal inputs alone, as well as their combination, can generate bursting. In all cases the impact of apical input is compared with the mode of apical isolation, in which there is no effect of apical input.

## 2 Methods

### 2.1 Model pyramidal cell

Simulations of probabilistic spike firing were run using the Bahl et al. (2012) reduced 20-compartment model of a layer 5 pyramidal cell. We used model 2 of Bahl et al. (2012), which can generate backpropagation-activated calcium (BAC) spike firing. Model code for the NEURON simulator (*neuron*.*yale*.*edu*) was obtained from ModelDB (*modeldb*.*science*). The compartmental structure and active biophysics of the model are outlined in Table 1. A cell schematic is shown in Figure 1. Full details of the parameter values are available in Bahl et al. (2012) and in the code itself.

**Table 1:** Structure and active biophysics of the Bahl et al. (2012) model. Cpts - number of numerical compartments; L - length (*μm*); Diam - diameter (*μm*); Nat - transient sodium current; Kfast - fast potassium; Kslow - slow potassium; Nap - persistent sodium; Km - muscarinic potassium; HCN - hyperpolarization-activated cation; Cas - slow calcium; KCa - calcium-activated potassium; CP - calcium pump.

| Section | Cpts | L | Diam | Nat | Kfast | Kslow | Nap | Km | HCN | Cas | KCa | CP |
| --- | --- | --- | --- | --- | --- | --- | --- | --- | --- | --- | --- | --- |
| soma | 1 | 23.1 | 23.1 | x | x | x | x | x |  |  |  |  |
| basal dendrite | 1 | 257 | 8.7 |  |  |  |  |  | x |  |  |  |
| apical dendrite | 5 | 500 | 5.9 | x | x | x |  |  | x |  |  |  |
| apical tuft | 2 | 499 | 6 | x | x | x |  |  | x | x | x | x |
| axon hillock | 5 | 20 | 3.5-2 | x |  |  |  |  |  |  |  |  |
| initial segment | 5 | 25 | 2-1.5 | x |  |  |  |  |  |  |  |  |
| axon | 1 | 500 | 1.5 |  |  |  |  |  |  |  |  |  |

**Figure 1.**
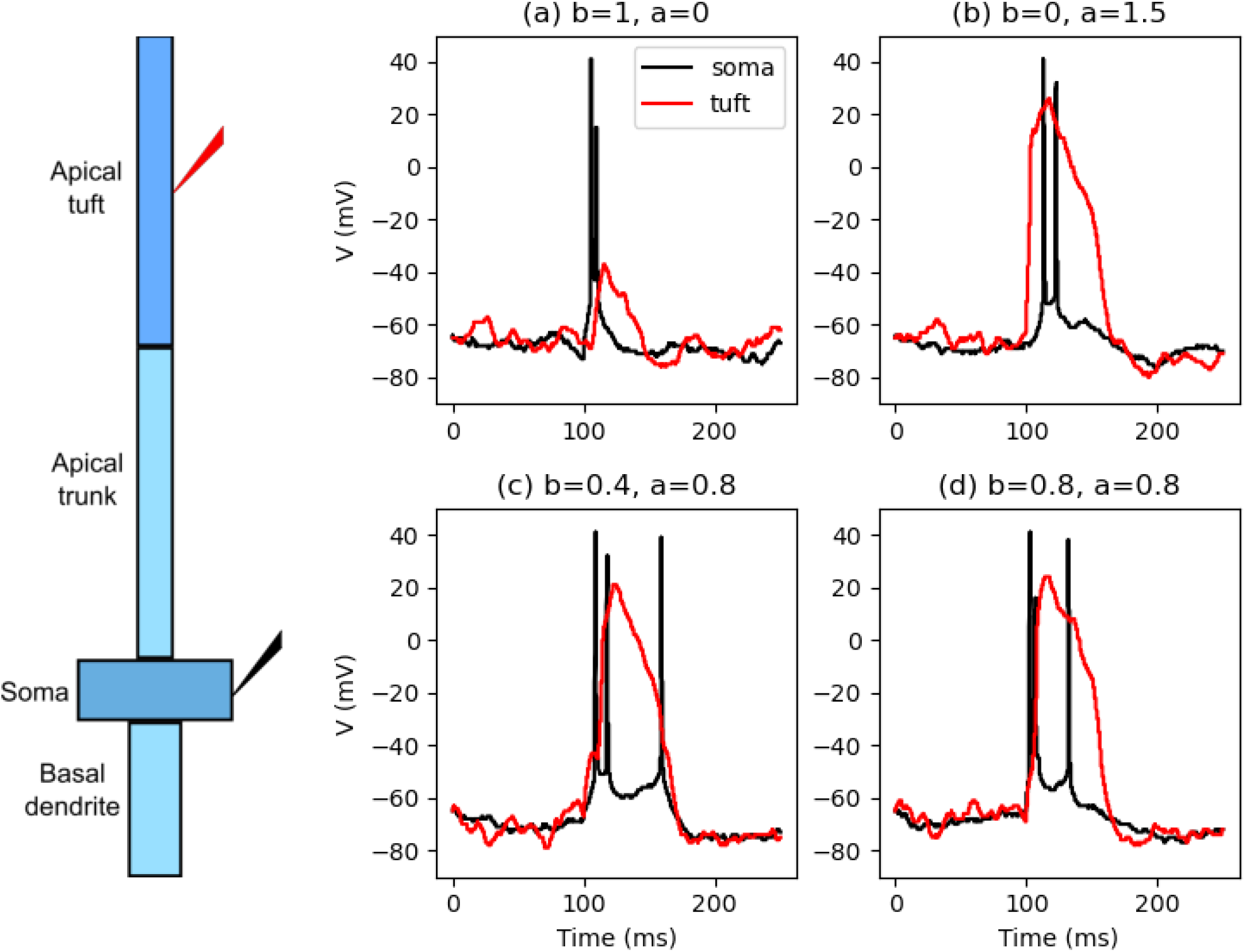
Examples of spiking responses to different stimulus strengths and durations. Cell schematic shows model structure and somatic (black) and apical tuft (red) stimulation and recording sites. For basal amplitude b nA and apical amplitude a nA and basal duration 10 ms: (a) basal input alone: b=1, a=0 (b) apical input alone: b=0, a=1.5; (c) low basal+apical: b=0.4, a=0.8; (d) BAC-firing: b=0.8, a=0.8.

#### 2.1 Model stimuli

Random background synaptic activity was modelled as a noisy current described by an Ornstein-Uhlenbeck (OU) process of the form used by Larkum et al. (2004). Our NEURON *nmodl* code was adapted from the synaptic conductance code of Destexhe et al. (2001). The current waveform is given by:

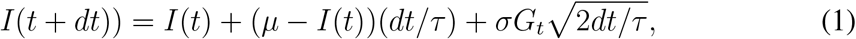

with mean current *μ* = 0 nA, standard deviation *σ* = 0.1 nA and correlation length *τ* = 3 ms. *G*_*t*_ is a Gaussian random number with mean 0 and a standard deviation of 1, chosen at each sample time *dt* = 25*μ*s. Independent noisy waveforms were injected into the middle of the soma and apical tuft sections.

The noisy currents themselves produced a very low probability of somatic spiking. Defined stimuli that could produce spiking were given simultaneously in the form of a short square-wave current pulse to the soma and an EPSP-like dual exponential current waveform to the apical tuft (rise time 0.5 ms; decay time 5 ms). Combinations of such stimuli of sufficient amplitude could produce BAC-firing, as recorded experimentally (Larkum et al., 1999, 2001; Larkum, 2013) and in pyramidal cell models (Bahl et al., 2012; Hay et al., 2011).

### 2.2 Simulations of stochastic spiking

Simulations of a single pyramidal cell with particular basal and apical stimulus amplitudes and time course were repeated 100 times with different noisy currents each time to generate a record of stochastic responses. Simulations were for 250 ms with the defined stimuli applied after 100 ms. Basal amplitudes were varied from 0 to 3 nA in 0.1 nA increments for step durations of 2 or 5 ms, and from 0 to 1 nA in 0.05 nA increments for a step duration of 10 ms. Apical amplitudes were varied from 0 to 1.7 nA in increments of 0.1 nA, always with an EPSP rise time of 0.5 ms and fall time of 5 ms. During each simulation, somatic spikes were counted, based on a somatic voltage threshold of -25 mV, from the onset of the defined stimuli to the end of the simulation. During analysis, groups of spikes recorded with interspike intervals of less than 25 ms were counted as bursts. Burst probability for given defined stimuli amplitudes was calculated as the fraction of the 100 simulations in which a burst occurred.

### 2.3 Information theoretic analysis

We consider a trivariate probabilistic system involving three discrete random variables: an output *Y* and two inputs *B* and *A*. Hence, underlying the discrete data sets we consider is a probability mass function Pr(*Y* = *y, B* = *b, A* = *a*), where *y, b, a* belong the the finite alphabets 𝒜_*y*_, 𝒜_*b*_, 𝒜_*a*_, respectively. Details of the information theoretic measures of such a system are given in Appendix A1.

The alphabets 𝒜_*b*_ and 𝒜_*a*_ consist of the sets of discrete basal and apical amplitudes used in the cell simulations. Each basal and apical input amplitude is treated as equally probable. The output spike count was categorised into two categories as 0-1 (no burst) and 1+ (burst) to allow calculation of burst probabilities.

In the analysis of the full data set with a basal duration of 10 ms, we consider basal and apical amplitudes in steps of 0.1 nA up to their range limits for each of four operating regimes (as defined in Section 3). The probability distributions for the four regimes are of size: 6 × 11 × 2 when basal and apical amplitudes are both low (range 0 to 1 nA for both); 6 × 18 × 2 when basal amplitude is low and apical amplitude is high (basal 0-0.5 nA; apical 0-1.7 nA); 11 × 11 × 2 for high basal amplitude and low apical amplitude (basal 0-1.0 nA; apical 0-1.0 nA); 11 × 18 × 2 when both amplitudes are high (basal 0-1.0 nA; apical 0-1.7 nA).

The probabilities in all cases are computed for each combination of basal input, apical input and spike count output as the number of occurrences of this combination divided by the product of the number of basal input amplitudes, the number of apical input amplitudes and 100.

#### Software for PID

The Ibroja PID (Bertschinger et al., 2014; Griffith and Koch, 2014) was estimated using compute UI (Banerjee et al., 2018) and the discrete information theory library dit (James et al., 2018). Python code was called from RStudio (R Core Team et al., 2021) by using the reticulate package (Ushey et al., 2020). The graphics were produced by using the ggplot2 package (Wickham, 2016) in RStudio.

### 2.4 Transfer function fitting

Burst probabilities were extracted from the simulation data in the form of burst frequencies over 100 repetitions at each basal and apical current amplitude. Bursts were counted as groups of spikes separated by interspike intervals of at most 25 ms. Derived transfer functions (see Appendix A2) were fit to this data by a least-squares fitting procedure using the SciPy (*scipy*.*org*) `least squares` function. Optimisation proceded simultaneously against all available basal and apical strengths for a particular duration of basal input (which was either 2, 5 or 10 ms). For each optimisation step, the burst probability as a function of basal amplitude when there was no apical input was calculated as *P*_2*b*_(*b*) (Equation 30). The resulting values were then used in the calculation of the full transfer function, *P*_2_(*b, a*) = *P*_1*b*_(*b*)[*P*_2*a*_(*a*)(1 − *P*_2*b*_(*b*)) + *P*_2*b*_(*b*)] (Equation 32), for all apical amplitudes greater than zero but in the low range, leading to the parameterisation of *P*_1*b*_(*b*) (Equation 29) and *P*_2*a*_(*a*) (Equation 31) and also allowing the calculation of the transfer function 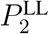 (Equation 2).

For apical input in the high range, the burst firing probability 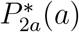 (Equation 35) was obtained by fitting to the full range of apical amplitudes when the basal input is zero. This function was then used to complete the transfer functions 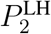 (Equation 4) and 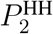 (Equation 5).

Standard errors in the parameter values were estimated from the Jacobian matrix and residuals. Fitting quality was also tested by refitting using weighted least squares and binomial nonlinear regression, both of which produced similar results.

## 3 Results

Computer simulations of a reduced-compartmental model of a thick-tufted layer 5 neocortical pyramidal cell (Bahl et al., 2012) are used to explore the contributions of basal and apical input streams to output burst firing. Details of the model and simulations are given in Section 2. In essence, through multiple independent simulations, we determine the likelihood that concomitant brief basal and apical inputs of known amplitude, on top of noisy background activity, cause an output burst. Output burst probability for these inputs is calculated as the fraction of simulations that result in a burst, and we calculate this over a range of input strengths. Information transmission of the two input streams is examined using partial information decomposition (PID) of this bursting data. We derive and fit transfer functions that give the probability of a burst as a function of the amplitudes of the basal and apical inputs.

### 3.1 Bursting regimes

Depending on the strength and duration of apical and basal inputs, a burst of 2 or more spikes separated by a short interspike interval can be generated in different ways: the apical or basal inputs alone may be strong enough to generate a burst, without the other input stream being active; weak apical and basal inputs, that alone cannot produce a burst or even a single output spike, may summate sufficiently to generate a burst; or basal input that causes a single output spike interacts with apical input through the back-propagation of this spike to then create a burst. This last case involves back-propagation activated calcium spiking in the apical dendrite, known a BAC-firing (Larkum, 2013). Examples of these different forms of burst generation are illustrated in Figure 1.

Multiple simulations were carried out across a range of current amplitudes for the basal and apical inputs to determine the probability of generating a burst with each combination of amplitudes. Figure 2 shows these results as contour maps of burst probability.

**Figure 2.**
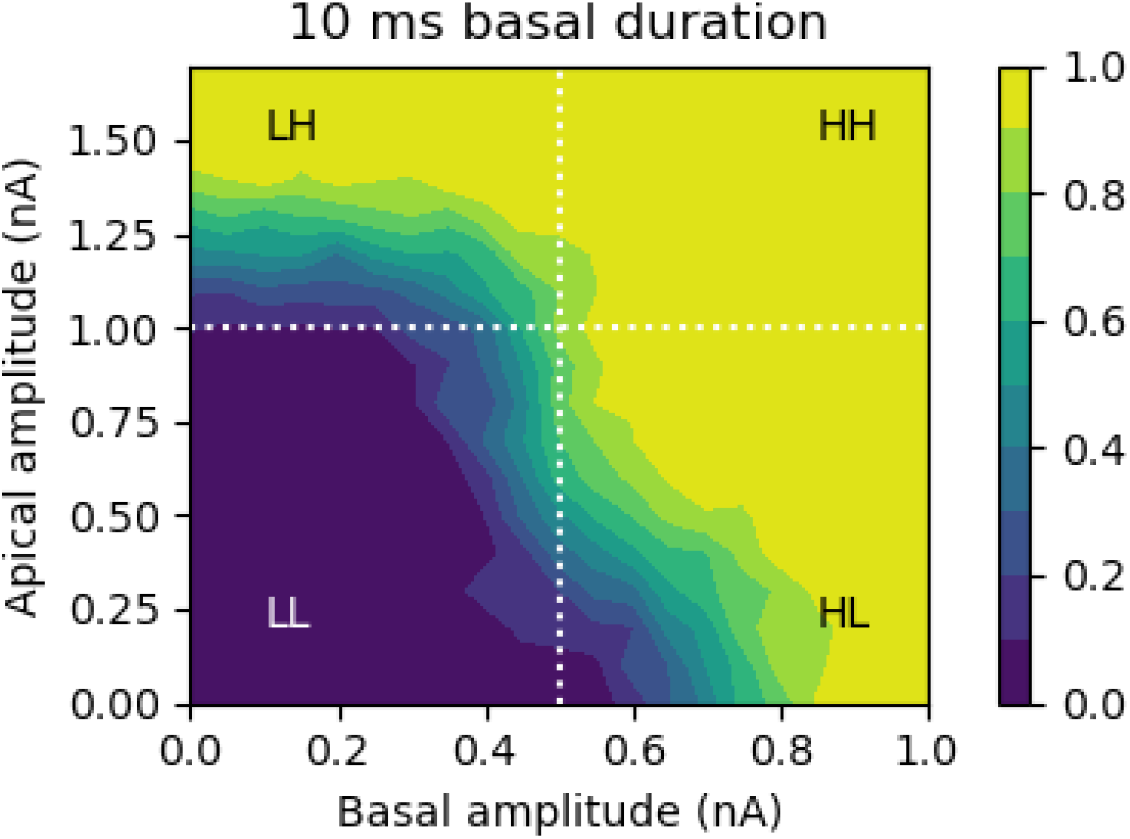
Contour map of burst probability for a basal duration of 10 ms, with basal (b) amplitude up to 1 nA and apical (a) amplitude up to 1.7 nA. Four operating regimes are indicated: (1) LL: *b* up to 0.5 nA, *a* up to 0.5 nA; (2) HL: *b* up to 1 nA, *a* up to 1 nA; (3) LH: *b* up to 0.5 nA, *a* up to 1.7 nA; (4) HH: *b* up to 1.0 nA, *a* up to 1.7 nA.

Taking as boundaries the basal and apical amplitudes at which their ability to produce a burst by themselves starts to increase from 0, this data can be divided into four different bursting regimes. Defining *low* ranges as being basal inputs up to 0.5 nA and apical inputs up to 1.0 nA, and *high* ranges as being basal inputs up to 1 nA and apical inputs up to 1.7 nA, gives the four regimes: (1) LL: *b=low, a=low*; (2) HL: *b=high, a=low*; (3) LH: *b=low, a=high*; (4) HH: *b=high, a=high*. These regimes are indicated in Figure 2; note that the high ranges subsume the low ranges. In the LL regime, only the highest values of basal and apical inputs in these ranges combined result in a significant probability of bursting, with neither input able to produce a burst by itself. In the HL regime the strongest basal inputs alone are able to generate a burst, but apical inputs in this range alone cannot. The LH regime reverses this so that strong apical inputs alone can produce a burst, but basal inputs cannot. The HH regime covers ranges of apical and basal inputs such that the strongest inputs of either pathway alone can produce a burst.

### 3.2 Information theory analysis of the bursting regimes

We apply information theoretic techniques to try to understand the information processing characteristics of these four identified bursting regimes. The question to be answered is what information does the occurrence of an output burst contain about the two distinct basal and apical input streams? Is information about each stream transmitted distinctly or is there only information about the combination of the inputs? Classical Shannon information theory sheds some light on these questions which are further illuminated by the use of partial information decomposition (PID). Details of these methods are given in Section 2 and Appendix A1.

#### Classical information measures

We first apply classical Shannon information theory to the burst probability distributions corresponding to each regime. Table 2 gives the values of of the classical information measures for these four regimes.

**Table 2:**
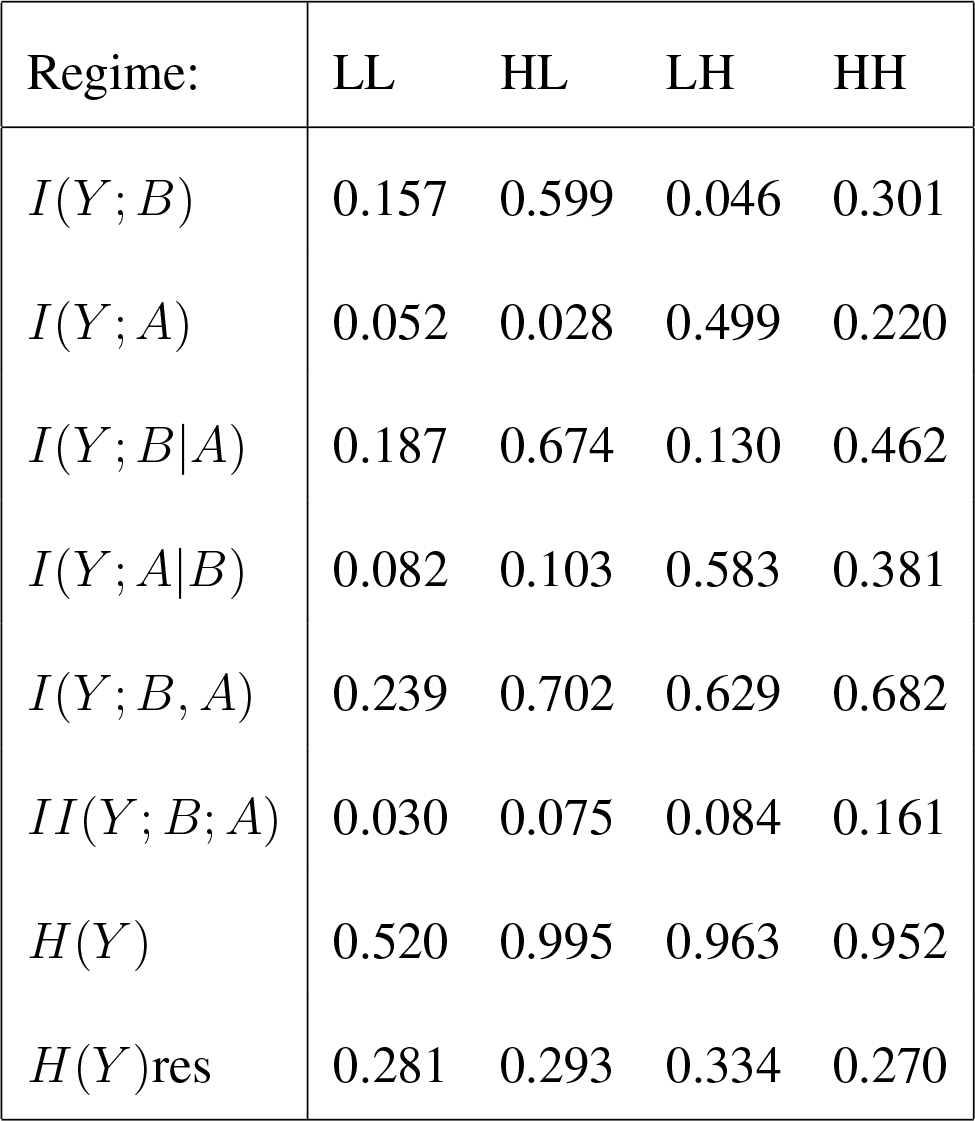
Classical information measures (bit) for four regimes of basal and apical inputs, with ‘*L*’ denoting ‘low’ and ‘*H*’ denoting ‘high’.

##### Regime LL: *b=low, a=low*

The joint mutual information, *I*(*Y* ; *B, A*), is low, at 0.239 bit, as is the value of the output entropy, at 0.520, so the transmitted information is only 46% of the output entropy, *H*(*Y*). The mutual information between each input and bursting output is low, but larger for the basal input, resulting in a unique information asymmetry in favour of the basal input of *I*(*Y* ; *B*|*A*) − *I*(*Y* ; *A*|*B*) = 0.105 bit. The value of the interaction information, *II*(*Y* ; *B*; *A*), is low but positive, indicative of some synergy between the three variables.

##### Regime HL: *b=high, a=low*

Adding higher levels of basal input leads to higher values for both the joint mutual information and the output entropy, with the transmitted information comprising a larger proportion of the output entropy (71%). The marginal mutual information between the basal input and bursting output, *I*(*Y* ; *B*), is now quite large (0.599), and is much larger than that between the apical input and bursting output (0.028). This means the unique information asymmetry in favour of the basal input is now substantial in magnitude and positive (0.571 bit). The value of the interaction information (0.075) is also positive and larger than in the LL regime, giving a larger lower bound for the synergy in the system.

##### Regime LH: *b=low, a=high*

Adding higher levels of apical input also results in higher values for both the joint mutual information and the output entropy than for the LL regime, with the transmitted information again comprising a large proportion of the output entropy (65%). However, the situation is reversed compared with the HL regime: it is now the marginal dependence between basal input and bursting output that is low (0.046 bit), being lower than in the LL regime, and much lower than the large value of 0.499 bit for the corresponding dependence between apical input and bursting output. The unique information asymmetry is again substantial in magnitude but now negative (−0.453 bit) and so in favour of the apical input. The value of the interaction information is again larger than for LL and similar to HL, giving a larger lower bound for the synergy in the system.

##### Regime HH: *b=high, a=high*

In the previous comparisons it is found that introducing higher levels of apical or basal input separately to expand the LL regime leads to a large value of unique information asymmetry, but what happens when higher levels of apical and basal input are added in combination? We find that the unique information asymmetry is much reduced in magnitude (0.081 bit in favour of the basal input) and that the marginal mutual informations between basal input and bursting output (0.301 bit), and between apical output and bursting output (0.22 bit), are similar in magnitude.

In summary, these information theoretic measures indicate that in the LL and HH regimes, both the basal and apical inputs contribute similarly to the information content in the output, whereas in the HL and LH regimes, the stronger input stream contributes significant information unique to itself, with little unique information conveyed about the weaker stream (less than in the LL and HH regimes).

#### Partial information decomposition

Partial information decomposition (PID) gives further insight and quantitative measures of the contributions of the two input streams to information in the output. This includes information unique to either input stream, shared between the streams, or synergistic, which is information that is not present in either input stream alone. Figure 3 shows a PID of the four operating regimes. The major sources of information contained in the bursting output are quite distinct in the different regimes.

**Figure 3.**
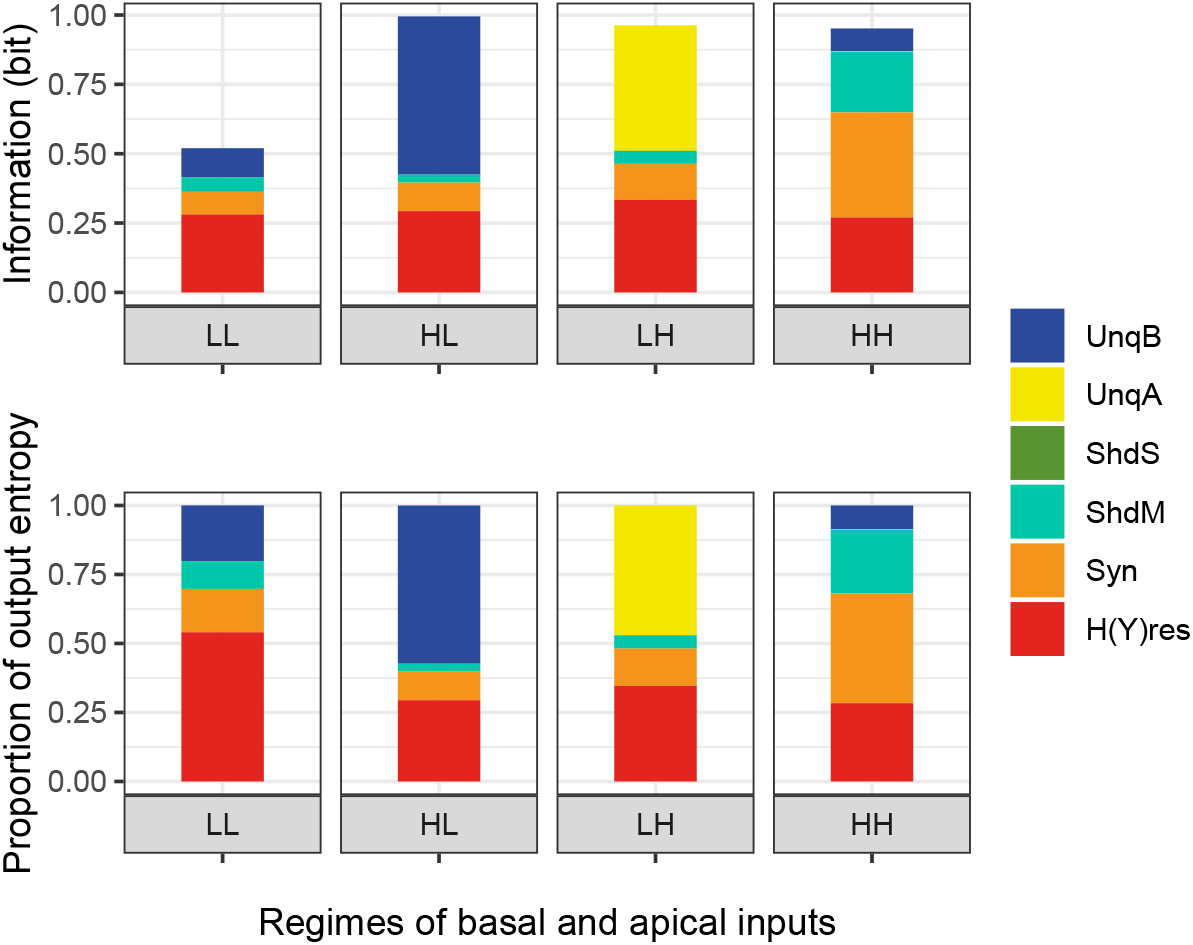
Absolute (top) and normalised (bottom) partial information decompositions obtained by using the Ibroja method (Bertschinger et al., 2014; Griffith and Koch, 2014) for each of the four regimes of basal and apical inputs. ‘L’ is an abbreviation of ‘low’, and ‘H’ is an abbreviation of ‘high’, so for example, ‘HL’ denotes the regime in which basal input is high and apical input is low.

##### Regime LL: *b=low, a=low*

Small amounts of information are transmitted that are unique to the basal input and due to mechanistic shared information and synergy, but information that is unique to the apical input is negligible.

##### Regime HL: *b=high, a=low*

The introduction of higher levels of basal input results in the transmission of more information, which is dominated by information unique to the basal input, with some mechanistic shared information and a slightly larger amount of synergy. Again there is negligible information unique to the apical input. However, since the apical input combines with the basal input in the transmission of mechanistic shared information and synergy, it does contribute to the output information about the basal input and thus the apical amplification of basal input is indicated.

##### Regime LH: *b=low, a=high*

The introduction of higher levels of apical input, but with low basal input, again results in the transmission of more information than in the LL regime, but it is now dominated by information unique to the apical input, with some mechanistic shared information and a slightly larger amount of synergy. There is now negligible information unique to the basal input. Since the basal input combines with the apical input in the transmission of mechanistic shared information and synergy the basal amplification of apical input is indicated.

##### Regime HH: *b=high, a=high*

Adding higher levels of both basal and apical input provides a dramatic change to the information decomposition. The unique components become very small, with the output being largely composed of synergy, and to a lesser extent mechanistic shared information, with a much smaller component unique to the basal input.

It is clear from this PID that the four operating regimes correspond to distinct modes of information processing. The HL regime has previously been termed *apical amplification* (Phillips, 2017, 2023) and corresponds with contextually-modulated information processing in which the apical inputs provide context which can amplify the response to the driving feedforward basal inputs. Conversely, the LH regime has been termed *apical drive* (Aru et al., 2020a; Phillips, 2023) and now the output is largely determined by internal sources which may be boosted by feedforward sensory evidence. The LL regime we choose to call a mode of *apical cooperation* as a large proportion of the output information is determined by both input streams combined. Finally, the HH regime we choose to call a mode of *apical integration* in which both input streams separately and in combination contribute to output information. This regime could therefore be described as being predominantly a generative mode of information processing in which its output depends upon both diverse internal sources, provided by the apical inputs, and more specific feedforward sources, provided by the basal inputs. Looking across these regimes, there is a key asymmetry shown by these results in that information unique to the apical input is transmitted only when apical input is high and basal is low (LH), whereas that unique to basal input is transmitted in all the other three regimes.

### 3.3 Transfer functions for the bursting regimes

In order to build a functional picture of how the two input streams combine in the different bursting regimes, we have constructed dual-input-stream transfer functions for the spike bursting probability. Details of how these functions have been derived are given in Appendix A2. The four related transfer functions covering the different operating regimes are as follows:

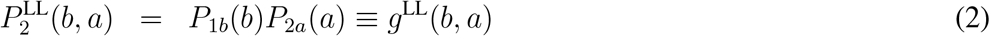

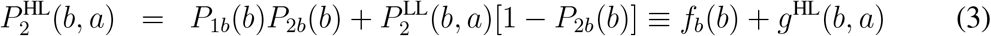

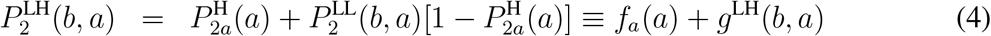

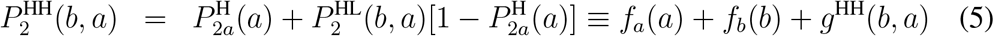

The functional breakdown in Equation 5 is obtained by expanding the second term to give *f*_*b*_(*b*) plus 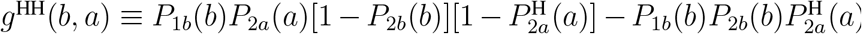.

These transfer functions are the combination of a number of distinct components (Equation 32) which can be interpreted as follows:

*P*_1*b*_(*b*) ≡ *P* (*Z*_1_ = 1|*b*) is the probability of an initial somatic spike (binary random variable *Z*_1_ = 1 when spike occurs) due to basal input alone.

*P*_2*b*_(*b*) ≡ *P* (*Z*_2_ = 1|*b*) is the burst probability (binary random variable *Z*_2_ = 1 when second spike (burst) occurs) due to basal input alone.

*P*_2*a*_(*a*) ≡ *P* (*Z*_2_ = 1|*Z*_1_ = 1, *a*) is the contribution of apical input to a full (or partial) calcium spike which can lead to a second (or more) somatic spike, following an initial somatic spike.

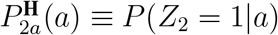 is the probability that apical input produces a calcium spike on its own.

*P*_1*b*_(*b*), *P*_2*a*_(*a*), *P*_2*b*_(*b*) and 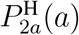 are sigmoidal functions of either basal stimulus amplitude, *b*, or apical stimulus amplitude, *a* (see the Appendix).

#### Modes of operation

The breakdown of these transfer functions shows explicitly how the apical and basal inputs combine in the different operating regimes and so provide distinct contributions to the information contained in the bursting output, corresponding with the four modes of information processing identified above. To further appreciate the distinct characteristics of these transfer functions, contour plots of their output across the relevant ranges of basal and apical input amplitude are shown in Figure 4. These can be compared with the simulation results shown in Figure 2.

**Figure 4.**
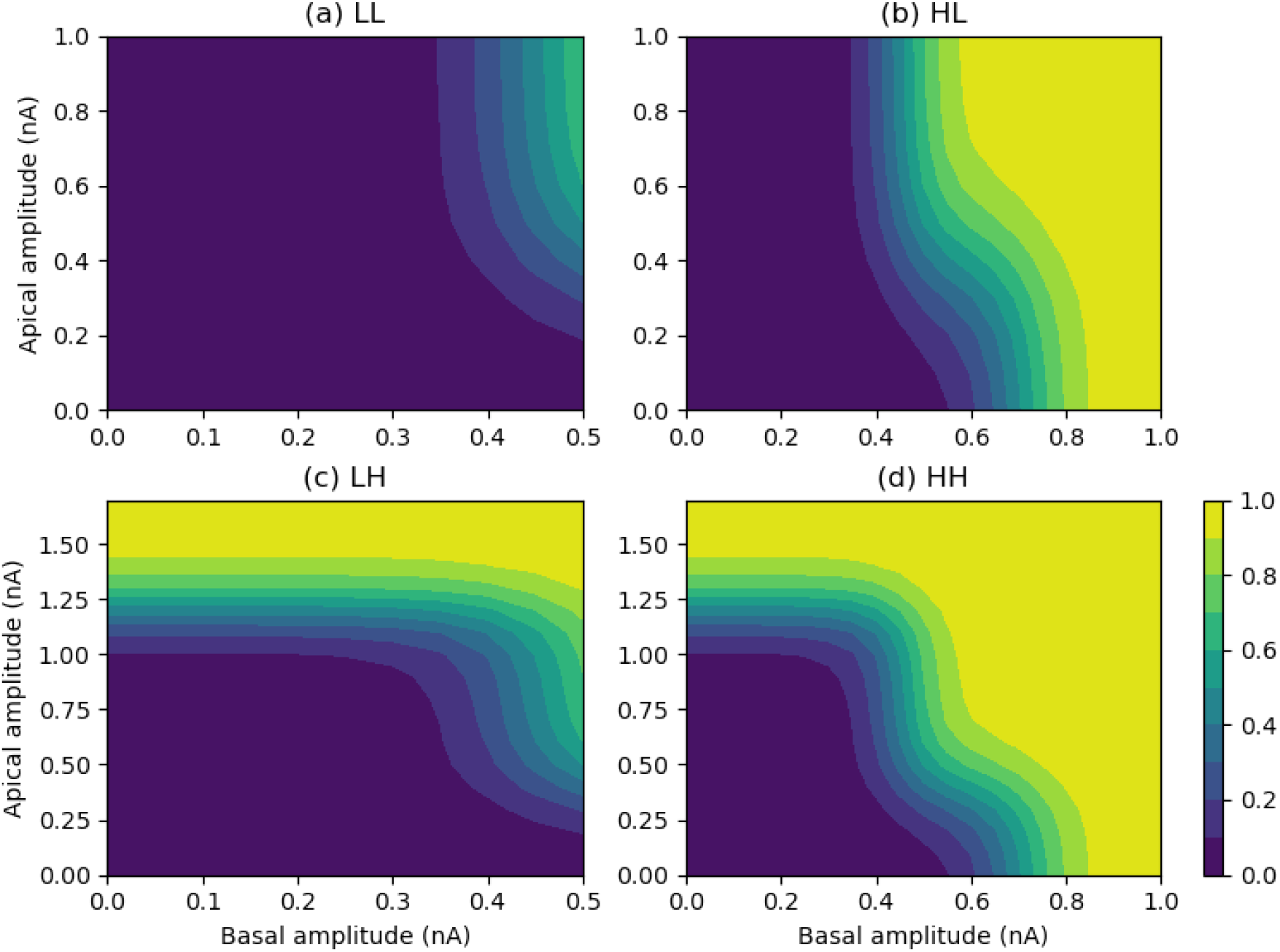
Contour plots of burst probability from the four transfer functions for different operating regimes for burst firing. (a) LL: basal=low, apical=low; (b) HL: basal=high, apical=low; (c) LH: basal=low, apical=high; (d) HH: basal=high, apical=high.

For basal and apical inputs both in their low ranges (LL: *apical cooperation*), bursting probability is clearly a function of both the basal and apical amplitudes combined (Figure 4a), as made explicit in Equation 2. This is not a simple summing, as bursting typically results from BAC-firing in which basal input triggers a single somatic spike which in turn combines with apical input to trigger a dendritic calcium spike and subsequent burst output.

With basal input in its high range, but apical input low (HL: *apical amplification*), bursting probability is strongly a function of basal amplitude, but with the transition from low to high bursting probability occurring at lower basal amplitudes with increasing apical amplitude (Figure 4b). This is captured by Equation 3 and is a functional expression of contextually-modulated information processing.

With low range basal input but high range apical input (LH: *apical drive*), bursting probability is almost purely a function of apical amplitude, with increased sensitivity in the upper reaches of the basal low range where the basal input starts to trigger single somatic spikes, which then starts to amplify the apical response (Figure 4c), as indicated by Equation 4.

For both inputs in their high range (HH: *apical integration*), bursting probability is a function of both the individual amplitudes and their combination, which is not a simple summation of the inputs, as indicated above for the other regimes, which this range subsumes. This is captured by Equation 5 and illustrated in Figure 4d. Close examination of Figure 4d shows how it combines the three other modes of operation. Purely apical drive is evident when the basal amplitude is below around 0.3. For low values of the apical amplitude (below around 0.1) burst probability is purely a function of basal amplitude, corresponding with what has been termed *apical isolation* (Aru et al., 2020a; Phillips, 2023). The interaction of basal and apical amplitudes is clear between these two extremes, encompassing both apical cooperation and apical amplification.

### 3.4 Examples of contextual information processing

#### Orientation selectivity

To assess the functional implications of these transfer functions, we consider the orientation selectivity example studied by Shai et al. (2015). This serves as a suitably general application for examining in-context and out-of-context processing, here meaning strong apical input either coincides with basal input or is out-of-sync with basal input. Orientation-specific basal and apical input stimuli are generated from von Mises circular distributions (Figure 5a,b). These stimuli are fed through the transfer functions to give the distribution of output bursting probability. Figure 5 illustrates the orientation selectivity of the transfer functions for the different operating regimes (Equations 2-5). In each example, the amplitudes of maximum basal and apical input, corresponding to peak orientation preference, are selected to fall within a prescribed operating regime, but near the boundaries of each regime so that there is a significant difference in the output response to basal and apical inputs alone compared with when they are combined. The left-hand panels show the output response when the basal and apical orientation preferences coincide at 0 radians, whilst the right-hand panels show the response when the apical preference is shifted to 2.5 radians.

**Figure 5.**
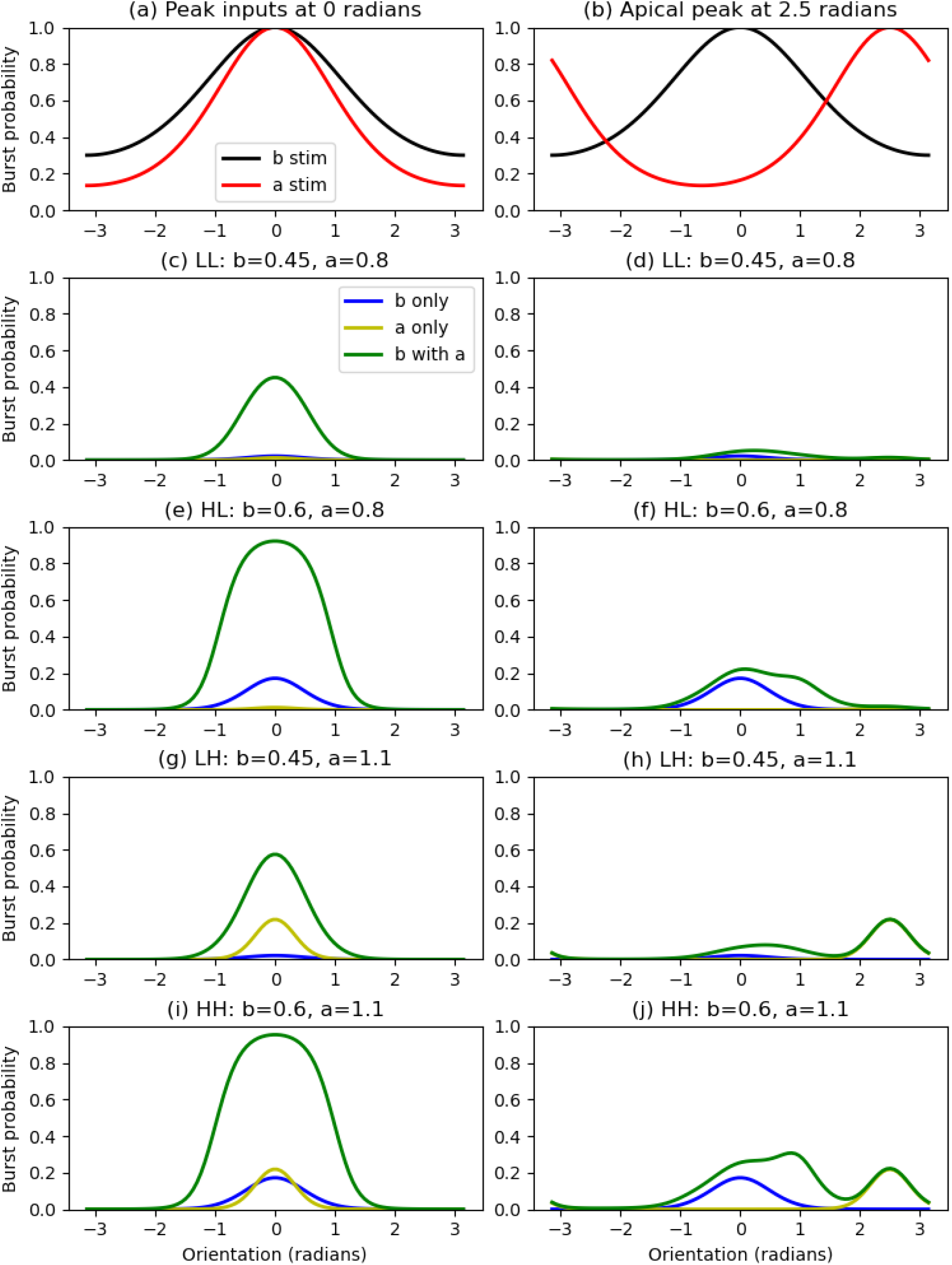
Orientation selectivity example of transfer function processing. Left-hand panels: both basal and apical inputs are tuned to an orientation of 0 radians; Right-hand panels: apical input now tuned to an orientation of 2.5 radians. Four operating regimes with peak basal b and apical a amplitudes: (c,d) LL: b=0.45, a=0.8; (e,f) HL: b=0.6, a=0.8; (g,h) LH: b=0.45, a=1.1; (i,j) HH: b=0.6, a=1.1.

In the LL regime (Figure 5c,d), peak responses to basal and apical inputs alone are small, but are greatly amplified when peak basal and apical input amplitudes coincide (Figure 5c). This amplification is much reduced when peak basal and apical amplitudes occur for different orientations and the peak bursting probability now occurs for an orientation between the preferred input orientations (Figure 5d).

The responses are similar in the HL regime (Figure 5e,f), but with the significant difference that when peak basal and apical amplitudes occur for different orientations, the peak bursting probability occurs at the preferred orientation of the basal input (Figure 5f). This is a clear example of contextually-modulated processing, where the contextual signal (the apical input) modulates the strength but not the selectivity of the driving signal (the basal input).

The impact of the basal and apical inputs is reversed in the LH regime (Figure 5g,h), with the basal input providing modest amplification of the response when orientation preferences coincide (Figure 5g) but having limited effect when the apical preference is shifted, with the peak response now occuring at the peak apical orientation (Figure 5h). The HH regime (Figure 5i,j) sees strong amplification when modest basal and apical inputs are combined with the same orientation preference (Figure 5i), but a weaker and rather broad response when the apical preference is shifted (Figure 5j). Now the peak output response occurs in between the preferred orientations of the inputs. This is clearly not a regime that provides tight orientation selectivity but rather averages the selectivity of its two input streams.

#### Lateral amplification and burst readout

To give a simple example of apical amplification and readout of bursts in a network of spiking cells, we simulated a purely excitatory 2-layer network of Bahl model pyramidal cells. In the input layer, each cell feeds lateral excitation to all of its neighbours via synapses in the apical tuft of each cell. This is loosely analogous to the long-range lateral intraregional connections in visual cortex that underlie contour integration, for example. For simplicity, shorter range intraregional inhibitory connections underlying lateral inhibition and normalization are ignored. Each input cell also sends a feedforward connection to a single readout cell via synapses with stochastic transmitter release. A schematic of this network is shown in Figure 6.

**Figure 6.**
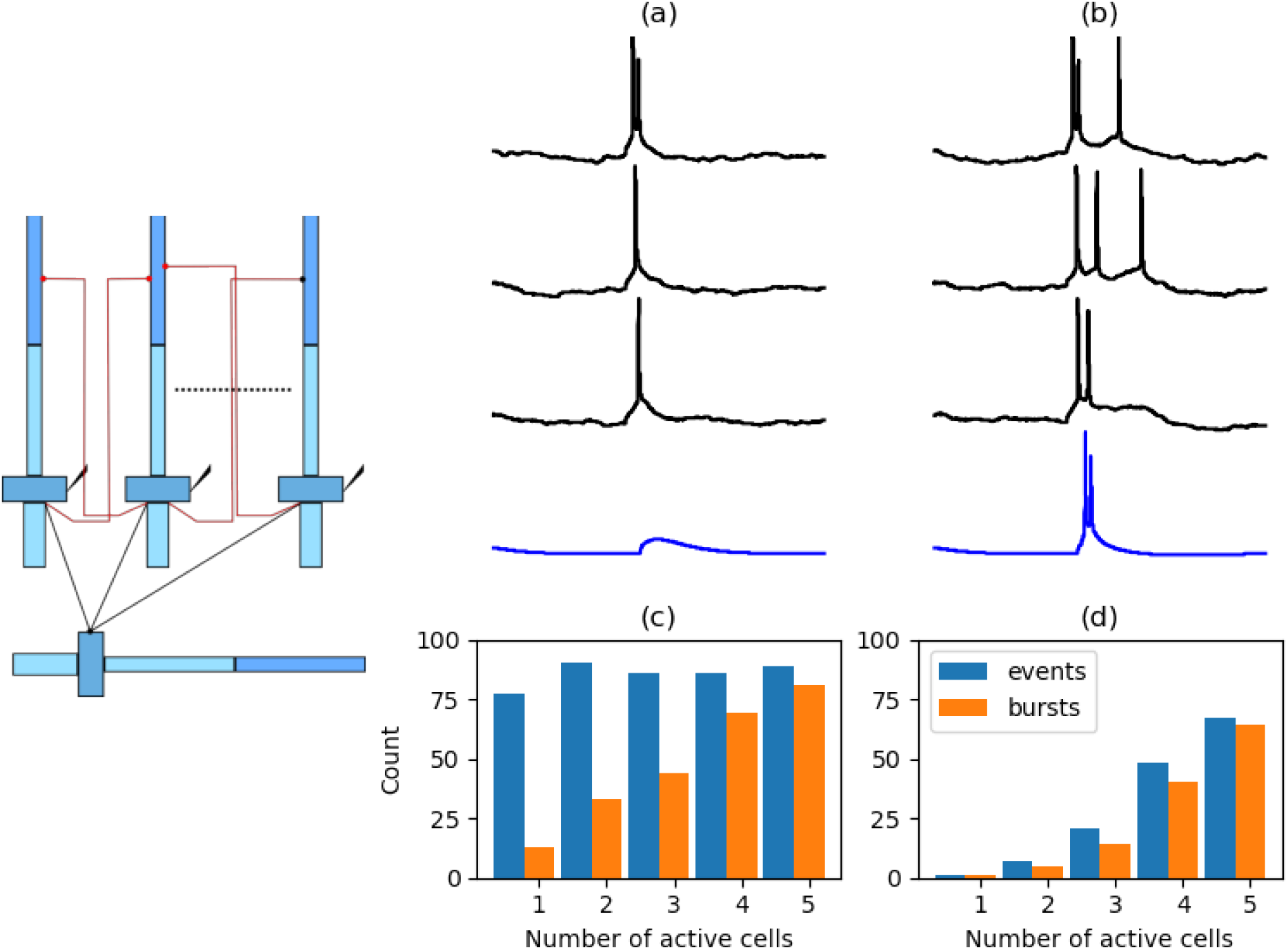
Lateral amplification and burst readout in a 2-layer network of spiking cells (schematic on left showing 3 of 5 input cells and the single output cell; see text for further details). (a) Somatic membrane potential in 3 example input cells (black) and the output cell (blue) when the input cells are not recurrently connected but are receiving basal current stimulation. (b) As for (a) but with the input cells having recurrent apical connections. (c) Response of the first input cell over 100 trials in the recurrently-connected network when from 1 to 5 of the input cells receive current stimulation (always including the first cell); the count of events include single spikes and bursts. (d) Response of the output cell over the same trials as in (c).

For the results shown here, the input layer consists of 5 cells, with each cell receiving basal stimulation via an injected current (duration 10 ms, amplitude 0.6 nA), and with the outputs of each cell feeding as a recurrent excitatory synaptic input (conductance rise time 0.5 ms, decay time 5 ms, amplitude 5 nS, reversal potential 0 mV, delay 2 ms) to the apical tuft of all of the other cells in this layer. The output layer consists of a single cell that receives feedforward somatic excitatory synaptic input (as for the recurrent synapses, but with a decay time of 10 ms) from each cell in the input layer. These synapses are stochastic so that an EPSP is only generated for a fraction of incoming action potentials (response probability is 0.2). This means a burst input is most likely to result in at least one EPSP and the output cell is most likely to spike itself when it is receiving bursting inputs.

When the input layer cells are not connected by recurrent apical synapses, the current stimulation to the input layer cells is sufficiently strong that they usually generate a single output spike in response, with an occassional burst (Figure 6a). In this case the output cell rarely responds with a spike of its own. However, when the input layer cells are connected by apical recurrent synapses, then the single spike is often converted into a burst due to initiation of a dendritic calcium spike (Figure 6b). This is apical amplification. Now the output cell often also responds with a spiking output as sufficient EPSPs occur through the stochastic synapses. Thus this cell is acting as a burst detector and is only responding to contextually-modulated activity (apical amplification through recurrent connections) in the input layer.

To further illustrate this contextual processing, 100 simulations were performed of the network with intact recurrent connections and with from 1 to 5 of the input cells receiving basal current stimulation. Figure 6c shows the spiking activity of the first cell, which is always one of the cells receiving current stimulation. When it is the only cell being stimulated it usually only produces a single spike as output, with occassional bursts. As the number of input cells being stimulated increases, then so does the apical input (context) of this cell and consequently the frequency of it generating a bursting output increases. The output cell rarely responds when only small numbers of input cells are activated (Figure 6d), but its own spiking output increases markedly as the input layer begins to generate a significant number of bursts (4 or 5 input cells being stimulated).

This demonstrates that laterally connected cells can act to amplify their neighbours’ responses when they respond to the same basal stimulus with an initial somatic spike. Note that such amplification only occurs when the lateral connections are via the apical tuft dendrites and not via the soma. Somatic connections do not add to the spiking response, at least for the short synaptic delays expected with nearby monosynaptic connections, as the input from a neighbour arrives within the refractory period following the spike originated by the basal stimulus.

It is sufficient for synaptic connections to have a low release probability, which is typical in cortex, for the receiving cell to effectively act as a burst-detecting filter. In our example this results in the read-out cell acting as a sensitive detector of contextually-modulated signal streams.

### 3.5 Comparison of transfer functions with two input streams

We have concentrated particularly on the HL regime in which the cell exhibits BAC firing (Larkum, 2013) leading to apical amplification (AA) of the output response. The resultant transfer function can be written so that it is explicity the sum of a function of the basal amplitude alone summed with an amplifying component that depends on both basal and apical input amplitudes, but is zero when the basal input is zero (Equation 3). This is the form of transfer function that has been proposed as the basis for contextually-modulated information processing (Kay et al., 2017; Kay and Phillips, 2020) in which the activation of a computational unit (neuron) due to receptive field (driving) inputs is modulated by a separate stream of contextual inputs. A basic transfer function with these characteristics is of the form (Phillips et al., 1995):

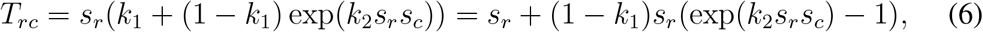

where *s*_*r*_ and *s*_*c*_ are the accumulated (weighted and summed) receptive field and contextual inputs, respectively. This is of the form *f* (*x*) + *g*(*x, y*) where *f* (*x*) = *x* and *g*(*x, y*) = *x* + (1 − *k*_1_)*x*(exp(*k*_2_*xy*) − 1). This is then be passed through a logistic function to give an output limited to between 0 and 1:

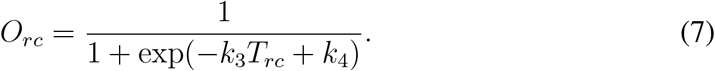

The use of computational units (neurons) with this transfer function in artificial neural networks has been demonstrated in a wide range of applications where the networks were trained to achieve contextually-modulated information processing using the *Coherent Infomax (CI)* learning rule derived using information theory (Phillips et al., 1995; Kay and Phillips, 1997; Kay et al., 1998; Kay and Phillips, 2011).

Using arguments based on consideration of the log odds for the generation of a second somatic spike, this basic transfer function has been extended to cover the situation of BAC firing in a pyramidal cell, as with our HL transfer function (Kay et al., 2019). The resultant function is:

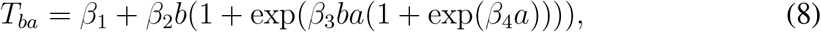

where *b* and *a* are the strengths of the basal and apical inputs, respectively. The main difference from the basic function (Equation 6) is that the contextual term includes the transformation of apical input into BAC firing via:

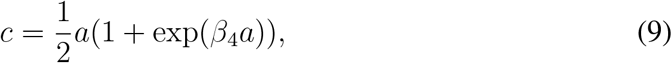

plus the addition of a constant *β*_1_ accounting for the prior odds of a burst.

Least-squares fitting to our simulated bursting data of the *T*_*ba*_ function (Equation 8), passed through a logistic (Equation 7) to give bursting probability, reveals that it has the same general characteristics of our more complex 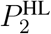 function (Equation 3) and gives a tolerable fit to the 10 ms basal data (Figure 7a), though not as good as our more complex function (Figure 7b).

**Figure 7.**
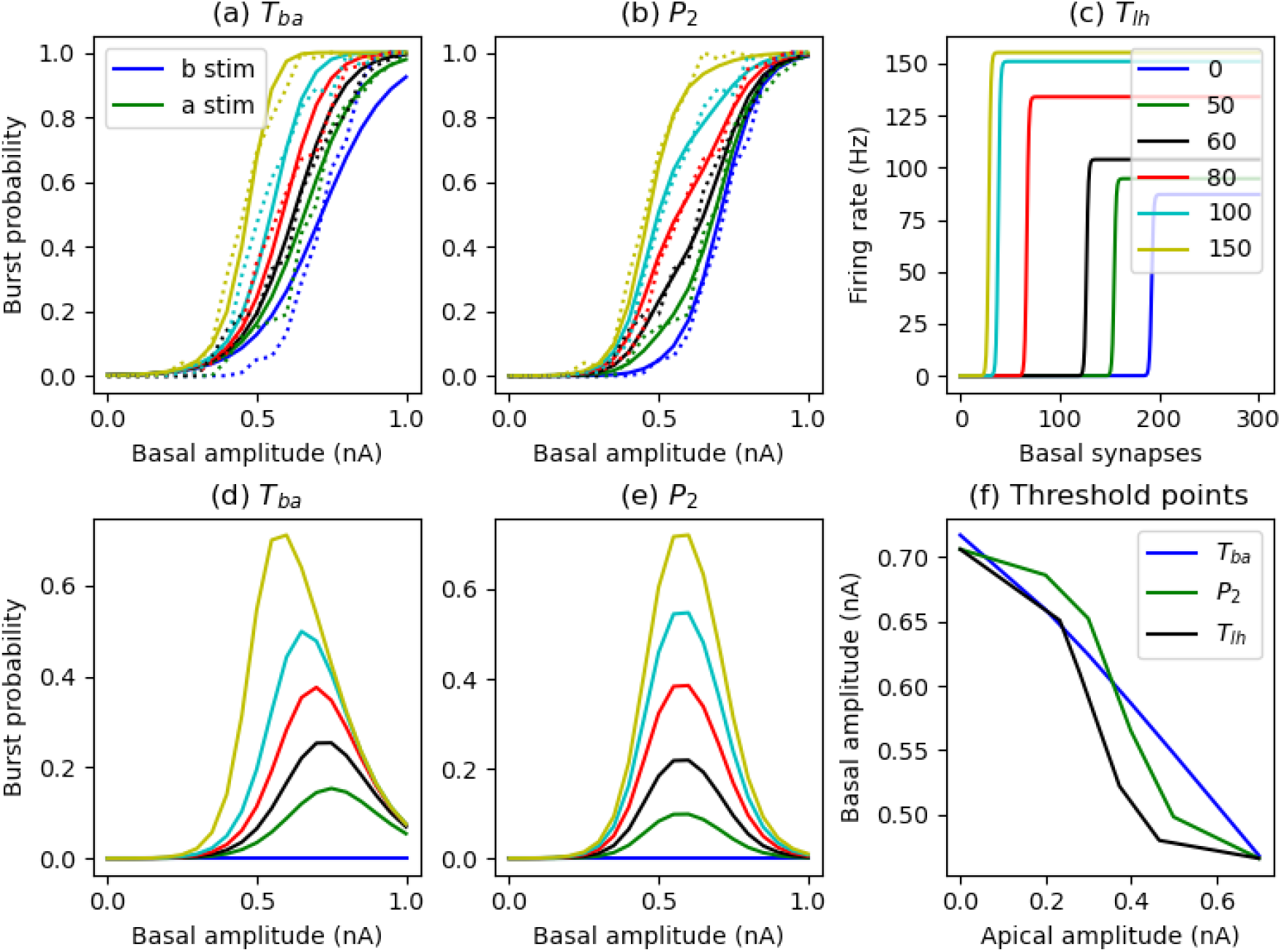
(a) *T*_*ba*_ transfer function through logistic (solid lines) with parameter values from least-squares fit to our 10 ms basal duration data, illustrated as burst probability versus basal amplitude, with plots for selected values of the apical amplitude (legend); (b) as for (a) but with our 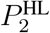 transfer function (note that dashed lines in (a) and (b) are the frequency data from simulations for a basal duration of 10 ms); (c) Shai et al. (2015) transfer function *T*_*lh*_ for firing frequency as a function of the number of basal and apical synaptic inputs; (d,e) difference between burst probability curves with apical input and burst probability with basal input alone, for the *T*_*ba*_ and 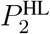 transfer functions, respectively; (f) combinations of apical and basal amplitudes required to achieve a threshold of 0.5 burst probability for the three transfer functions (note that the numbers of synaptic inputs for *T*_*lh*_ have been scaled to match the input currents of 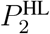 for comparison purposes).

The effect of apical input in the *T*_*ba*_ function is to increase the gain of the bursting response to basal input. In our 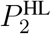 function, the response is left-shifted with increasing apical input, but by different amounts over the basal range, with low bursting probabilities being more quickly enhanced by apical input than higher probabilities. For 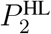, the response curves for no apical input and for strong apical input are essentially identical in shape and hence gain, but over different basal ranges. In between, the gain is effectively lowered, resulting in a greater dynamic range in the response to basal inputs (see particularly the response when apical input is 0.4 nA in Figure 7b). By considering the difference between response curves with non-zero apical input and the response with only basal input, it can be seen that apical amplification is particularly effective over a limited basal range for both the *T*_*ba*_ and 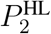 functions (Figure 7d,e). The point of greatest sensitivity to apical input shifts to lower basal values for *T*_*ba*_, but for 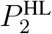 (and hence the pyramidal cell) it is constant at around a basal amplitude of 0.6 nA.

In a closely related study, Shai et al. (2015) used simulations of a detailed compartmental model of a layer 5 pyramidal cell to examine its firing rate response to different numbers of concomitant basal and apical synaptic inputs. Though not classifying the output as bursting or not, they demonstrated that the cell exhibited a rapid transition between low and high firing rates, as calculated over a 100 ms period. The high firing rates had a strong dependence on calcium levels underpinning BAC firing. They collected data across a range of numbers of basal and apical synaptic inputs, but with only a single simulation per pair of numbers, resulting in hard thresholds between low and high firing. They fitted a transfer function to this data in which output firing is driven by the basal input, but where the maximum firing rate *M* and the threshold *T* between low and high firing, were both functions of the number of apical inputs:

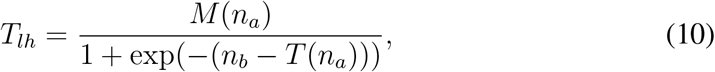

where *n*_*a*_ and *n*_*b*_ are the number of apical and basal synaptic inputs. The maximum rate *M* (*n*_*a*_) is an increasing sigmoidal function of *n*_*a*_, whilst the threshold *T* (*n*_*a*_) is a decreasing sigmoidal function of *n*_*a*_. This function, as fit by Shai et al. (2015) to their simulated data, is shown in Figure 7c.

A similar approach is taken by Pastorelli et al. (2023) where they fit a piece-wise linear transfer function to spiking data from a reduced two-compartment pyramidal cell model. In their case, firing frequency due to somatic and dendritic injected step currents is measured over a long 2 second time period. They fit 2-dimensional planes to the distinct regions of high and low firing rate.

To compare our transfer function with that of Shai et al. (2015), we can define a threshold of 0.5 for the transition from a low to a high bursting probability. The paired apical and basal input amplitudes that achieve this threshold for the *T*_*ba*_ and 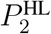 transfer functions are plotted in Figure 7f. In this same figure, equivalent threshold points (black line) for the *T*_*lh*_ function have been found by scaling the firing rate responses shown in Figure 7c to the maximum achieved in each case and then scaling the numbers of apical and basal synaptic inputs to the ranges of input currents of 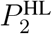. In this way it can be seen that the data and transfer functions of our model and that of (Shai et al., 2015) are related, with 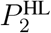 and *T*_*lh*_ showing a similar non-linear decrease in basal amplitude required to reach the threshold, with increasing apical amplitude. This decline is quite linear for *T*_*ba*_.

## 4 Discussion

We have examined input stream integration and subsequent information processing in one class of neocortical neurons, namely thick-tufted layer 5 pyramidal cells. These cells broadly have two anatomically-separated basal and apical input streams. These streams interact via the generation of a dendritic calcium spike to cause or amplify output burst firing. Though single spiking is also a unit of output currency, bursting is a particular indication of the interaction of the two input streams and we have derived transfer functions to capture this interaction. We now discuss this and related work on the information processing possibilities of two input streams and subsequent bursting outputs, and experimental evidence for the existence of the modes of operation we have identified.

### 4.1 Information processing with two input streams

The nature of the information processing carried out by neocortical pyramidal cells that receive anatomically-distinct basal (perisomatic) and apical dendritic inputs from different sources is the subject of many investigations and hypotheses (Payeur et al., 2019). Our study demonstrates the ability of pyramidal cells that exhibit BAC firing to carry out contextually-modulated processing of the sensory-derived feedforward basal inputs, where the distinct apical inputs provide the relevant context. Output bursts are the information carriers. A simple orientation-selectivity example illustrates how the appropriate context can greatly amplify the response to the sensory (feedforward) input. The out-of-context response is reduced but still has the same receptive field peak i.e. the conflicting context does not shift this peak.

Information theory has been used to understand what information our transfer functions transmit about the basal and apical input streams. Partial information decomposition reveals that in a regime where sufficiently strong basal input is able to generate a burst, but apical input alone is not, then the presence or not of a burst carries significant information about the strength of the basal input, but little about the apical input (Figure 3 HL). There is some shared and synergistic information. This has been termed the *apical amplification* processing mode (Phillips, 2017, 2023). The situation is reversed if the apical input is strong enough to cause a burst by itself, but the basal input is not (Figure 3 LH), which has been termed the *apical drive* mode (Aru et al., 2020a; Phillips, 2023). In the situation where neither the basal nor apical inputs alone can generate a burst, the output contains significant shared and synergistic information, with some unique information about the basal strength as it contributes to bursting through the generation of an initial sodium spike, even though it cannot generate a burst by itself (Figure 3 LL). We denote this the *apical cooperation* mode. When both inputs are strong enough to generate bursting by themselves (HH) then most information is shared and synergistic, as the probability of bursting is a function of the nonlinear summation of the basal and apical inputs (Figure 3 HH). We denote this the *apical integration* mode.

In the *apical cooperation* mode (low input regime, LL), where the individual inputs do not generate bursts by themselves, Naud and Sprekeler (2018) demonstrate that single spike and bursting outputs can carry multiplexed information about the basal and apical drives to a pyramidal cell, with different downstream circuits able to readout these different signals. In layered networks, short-term depression in ascending basal connections allows the bottom-up spike rate to be transmitted, irrespective of bursting induced by top-down inputs. Short-term facilitation in descending dendritic connections, on the other hand, allows information on top-down inputs to be transmitted, in the form of burst probability or burst rate. This separation of signals relies on operating in a regime where basal inputs alone cannot produce bursts.

The *apical integration* mode opens up the possibility of new forms of information processing that have been termed *information modification* (Wibral et al., 2017a) or *coding with synergy* (Wibral et al., 2017b), in which new information is generated in the output that is not available in either of the input streams individually. We would conjecture that our mode of apical integration captures a mechanism underpinning the formation of thoughts and imagination in the awake animal (Phillips, 2023).

### 4.2 Two-stream signal interaction in cortical pyramidal cells

There is increasing *in vivo* experimental evidence as to the influence of contextual signals on the receptive field responses of cortical pyramidal cells (PCs). In the visual system, there is considerable evidence that PC receptive field responses can alter in task-dependent ways, indicating contextually-modulated processing (Gilbert and Li, 2013; Pakan et al., 2018). Experiments in primates show that neural responses to visual inputs in their receptive field can be altered by nearby flanking signals that may be relevant or irrelevant to the task at hand, enabling such tasks as contour integration (Gilbert and Li, 2013). Our simple network example of lateral amplification provides an indication of a possible mechanism underpinning such effects (Figure 6).

New experimental techniques are now allowing the cellular and network effects and mechanisms of contextual modulation to begin to be unpicked, particularly in the mouse visual system (Pakan et al., 2018). In rodents, attention and locomotion increase the response amplitude and selectivity of PC receptive fields to visual input (Pakan et al., 2018), such as orientation selectivity (Niell and Stryker, 2010). Firing rate increases during locomotion, compared to rest, have been measured in individual PCs, corresponding to both additive and multiplicative changes in receptive fields (Dadarlat and Stryker, 2017). Importantly, receptive field orientation preference is not changed, but the increased response amplitude increases orientation discriminability. This corresponds with the effects seen in the example of orientation selectivity with our identified transfer function (Figure 5). In cortex, such effects likely will be mediated by cellular and network-level effects including changes in inhibition and neuromodulation (Dadarlat and Stryker, 2017; Pakan et al., 2018), but also through the increased influence of contextual excitatory inputs to PCs on their spiking output. The combination of locomotion and visual input can be a better predictor of PC activity than visual input alone (Keller et al., 2012). Further, visual activity has been recorded due to locomotion in the dark, indicating that contextual (apical) drive can generate output spiking on its own, without corresponding feedforward sensory inputs (Keller et al., 2012).

Dendritic calcium spiking in layer 5 PCs is correlated with cognition in somatosensory cortex (Xu et al., 2012). More specifically, experiments in mice show that calcium spikes in the apical dendrites of layer 5 PCs affect the detection of whisker stimuli in barrel cortex, corresponding with a lowering of stimulus intensity threshold for detection (Takahashi et al., 2016). These experiments showed that blocking of calcium spikes shifts the detection threshold to the right (larger stimulus amplitude), indicating that dendritic spiking modulates cell output to lower the response threshold to sensory input. Further, upregulating excitability in dendrites leads to an increased false detection rate, giving evidence of apical drive. Somatic bursts do correlate with detection though they are less predictive of detection than the overall firing rate (Takahashi et al., 2016).

### 4.3 Control of two-stream signal interaction in cortical pyramidal cells

We have identified different operating regimes for layer 5 pyramidal cells that depend on the relative strengths of the excitatory basal and apical input streams. Which regime may be in operation in the behaving animal is determined not purely by the strengths of the excitatory inputs, but also is controlled dynamically by inhibition and neuromodulation. In cortical circuits, the interaction of apical and basal excitatory inputs is strictly controlled through inhibition, mediated by a variety of classes of interneuron that make layer-specific connections onto pyramidal cells and are preferentially driven by top-down, bottom-up or lateral connections (Gentet, 2012). Perisomatic inhibition controls output spiking, with the potential to limit bursting output despite the initiation of dendritic calcium spikes. On the other hand, dendritic calcium spikes are controlled by inhibitory interneurons that target the apical tuft dendrites, affecting calcium spike initiation, or the apical trunk, affecting their transmission to the soma (Murayama et al., 2009; Schulz et al., 2021). In these highly polarised cells, dendritic inhibition has little direct affect on output spiking driven by basal inputs. Leleo and Segev (2021) used detailed computer simulations of a layer 5 pyramidal cell to explore the spatio-temporal effects of inhibition on dendritic spiking and subsequent burst firing.

Excitatory pathways influence their own impact on target pyramidal cells through which classes of inhibitory interneuron they also connect with. Of particular interest is the disinhibition of apical dendrites that promotes the generation of calcium spikes, through interneurons that specifically target other interneurons that inhibit the apical tuft and trunk (Gentet, 2012; Jiang et al., 2013). This disinhibition is quite focal within neocortical columns (Jiang et al., 2013) and allows gating of interactions between the basal and apical inputs that can mediate contextually-modulated information transmission (Wang and Yang, 2018). It appears that cross-modal contextual inputs preferentially disinhibit their target pyramidal cells, whereas within the same modality such inputs largely inhibit their target PCs (Shen et al., 2022; Wang and Yang, 2018).

The influence of apical tuft activity on layer 5 pyramidal cell output strongly depends on the neuromodulatory state of the cell. High levels of acetylcholine and norepinephrine are present in the neocortex during attention and active behaviour in the awake animal. These neuromodulators regulate both pyramidal cell and interneuron activity. In particular, activation of muscarinic acetylcholine receptors in the apical tuft of layer 5 pyramidal cells promotes calcium spiking through upregulation of R-type calcium channels (Williams and Fletcher, 2019). Adrenergic modulation also increases tuft excitability, putatively through blocking of dendritic HCN channels (Labarrera et al., 2018; Mäki-Marttunen and Mäki-Marttunen, 2022). General anesthesia acts to decouple the apical tuft from the cell body, putatively through blocking metabotropic glumate and cholinergic receptors in the apical trunk that promote coupling in the awake animal (Phillips et al., 2018; Suzuki and Larkum, 2020).

The effect of apical input may be controlled in different ways that may seem similar from the cell output point of view, but have distinct effects within the cell. Inhibition and neuromodulation may limit the summed strength of apical tuft synaptic inputs, or the ability of these inputs to generate a large calcium spike, or the transmission of this spike to the cell body (Leleo and Segev, 2021; Pastorelli et al., 2023). All of these will limit the contribution of the apical input stream to bursting output. Dendritic inhibition of synaptic input is closest to the simple change in amplitude we have studied here. Limiting transmission of the calcium spike to the cell body, on the other hand, while still allowing a dendritic calcium spike, may be vital for enabling synaptic plasticity in the apical tuft, while preventing its contribution to cell output (Leleo and Segev, 2021). We have not considered synaptic plasticity here, but it is of fundamental importance to establishing information storage and transmission in which regenerative activity in the apical dendrites and bursting may play a significant part (Payeur et al., 2021; Francioni and Harnett, 2022; Baronig and Legenstein, 2024).

While neocortical pyramidal cells in general usually have the same basic arrangement of apical and basal inputs, thick-tufted layer 5 pyramidal cells have the most distinct nonlinear interaction between contextual apical and feedforward basal inputs mediated by apical dendritic calcium spikes. However, even within this class of cells, BAC firing and subsequent bursting is limited to the largest cells (Fletcher and Williams, 2019; Galloni et al., 2020). Increasing apical trunk length results in greater dendritic compartmentalisation and a greater propensity for generating large calcium spikes that are still able to promote burst firing at the soma. BAC firing is enabled by active, sodium-channel-mediated back propagation of somatic action potentials (Galloni et al., 2020). Shorter cells only exhibit limited calcium spikes that do not trigger bursts but apical inputs can generate single spike firing in the soma. Shorter cells are more electrically compact which is enhanced by a reduced axial resistance in the apical trunk (Fletcher and Williams, 2019). These differences correspond with cortical location and its corresponding thickness, with a rostral-caudal gradient from large to small in primary visual cortex (Fletcher and Williams, 2019) and a distinct difference between primary (larger cells) and secondary (smaller cells) visual areas (Galloni et al., 2020) in the rat. At the other extreme, human pyramidal cells are larger and consequently exhibit even greater compartmentalisation of the distal apical dendrites, to the extent that calcium spiking in the dendrites does not lead to bursting in the soma (Beaulieu-Laroche et al., 2018). Thus two-stream signal interaction in pyramidal cells takes a variety of forms depending on cortical area and animal species.

## Conclusion

Partial information decomposition and transfer function fitting have been used to characterise the input-output properties of burst firing in a stochastic model of a layer 5 neocortical pyramidal cell that can exhibit BAC firing (Bahl et al., 2012). It is revealed that the cell can operate in different information processing modes, depending on the amplitude ranges of the basal and apical input streams. Highlighting the contribution of the apical stream, these modes have been termed apical amplification, apical drive (Phillips, 2017; Aru et al., 2020a; Phillips, 2023), apical cooperation and apical integration. The encompassing theme of these modes is contextually-modulated information processing in which contextual apical inputs from diverse brain regions refine signal transmission of feedforward sensory inputs.

These different modes are plausibly obtained *in vivo* through the activation of targeted inhibitory pathways and network neuromodulation. Further modelling work can address this explicitly (Leleo and Segev, 2021; Mäki-Marttunen and Mäki-Marttunen, 2022; Pastorelli et al., 2023).

As indicated above, the nature of the interaction between basal and apical inputs is very much pyramidal cell-type specific. The approach we have taken here, using a combination of transfer function fitting and information theory can be used to characterise other classes of pyramidal cell and provide a comprehensive picture of contextually-modulated information processing across the neocortex.

## Appendix

### A1. Information theory notation and definitions

We consider a trivariate probabilistic system involving three discrete random variables: an output *Y* and two inputs *B* and *A*. Hence, underlying the discrete data sets we consider is a probability mass function Pr(*Y* = *y, B* = *b, A* = *a*), where *y, b, a* belong the the finite alphabets 𝒜 _*y*_, 𝒜 _*b*_, 𝒜 _*a*_, respectively.

#### 4.1 Classical information theory

We now define the standard information theoretic terms that are required in this work and they are based on results in (Cover and Thomas, 1991). We denote by the function *H* the usual Shannon entropy, and note that any term with zero probabilities makes no contribution to the sums involved. The joint mutual information that is shared by *Y* and the pair (*B, A*) is given by,

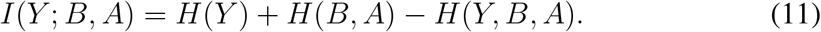

The information that is shared between *Y* and *B* but not with *A* is

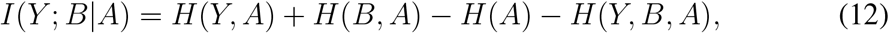

and the information that is shared between *Y* and *A* but not with *B* is

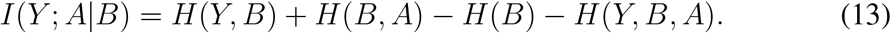

The information shared between *Y* and *B* is

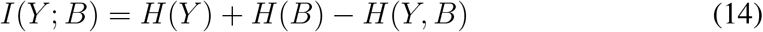

and between *Y* and *A* is

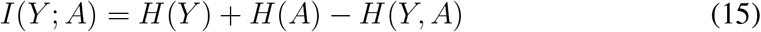

The interaction information (McGill, 1954) is a measure of information involving all three variables, *Y, B, A* and is defined by

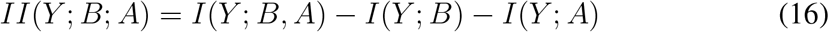

McGill’s interaction information has been used as a measure of synergy (Schneidman et al., 2003), with a positive value indicating the presence of synergy and a negative value indicating redundancy. See also (Gat and Tishby, 1998). The negative of McGill’s measure has been termed *coinformation* (Bell, 2003), and it has been used as an objective function in an artificial neural network possessing two distinct sites of integration (Phillips et al., 1995). In the case of three variables, the negative of McGill’s measure is a special case of the general O-information measure (Rosas et al., 2019). The O-information measure can be particularly useful with systems which have a large number of interacting variables (Varley et al., 2023).

#### 4.3 Partial Information Decomposition

The output entropy, *H*(*Y*), may be written as

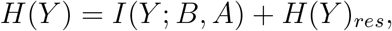

where *H*(*Y*)_*res*_ is the residual output entropy.

The decomposition of the joint mutual information can be expressed as (Wibral et al., 2017b):

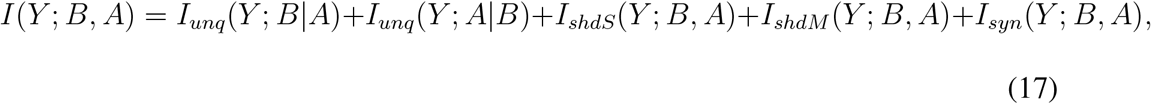

where the shared information *I*_*shd*_(*Y* ; *B, A*) has been split into two separate components of source shared or mechanistic shared information (Pica et al., 2017) as:

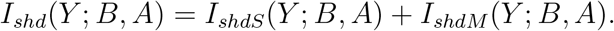

Adapting the notation of (Wibral et al., 2017b) we express our joint input mutual information in four terms as follows:

Unq*B* ≡ *I*_*unq*_(*Y* ; *B*|*A*) denotes the unique information that *B* conveys about *Y* ;

Unq*A* ≡ *I*_*unq*_(*Y* ; *A*|*B*) is the unique information that *A* conveys about *Y* ;

ShdS ≡ *I*_*shdS*_(*Y* ; *B, A*) gives the shared (or redundant) information that both *B* and *A* have about *Y*, due to the correlation between *A* and *B*. This is termed source shared information.

ShdM ≡ *I*_*shdM*_ (*Y* ; *B, A*) gives the shared (or redundant) information that both *B* and *A* have about *Y*, due to the probabilistic mechanism. This is termed mechanistic shared information.

Syn ≡ *I*_*syn*_(*Y* ; *B, A*) is the synergy or information that the joint variable (*B, A*) has about *Y* that cannot be obtained by observing *B* and *A* separately.

For an excellent tutorial on information theory and partial information decomposition, with illustrations from neuroscience, see (Timme and Lapish, 2018). It is possible to make deductions about a PID by using the following four equations which give a link between the components of a PID and certain classical Shannon measures of mutual information. The following are in (Wibral et al., 2017b) (eqs. 4, 5 with amended notation); see also (Williams and Beer, 2010).

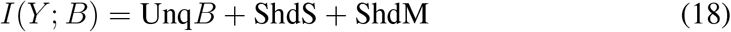

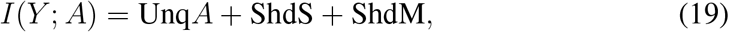

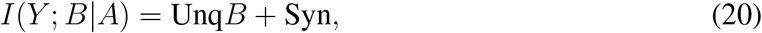

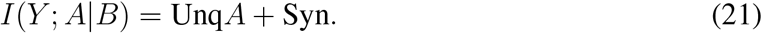

Using (17), (18), (19) we may deduce the following connections between classical information measures and partial information components:

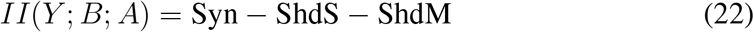

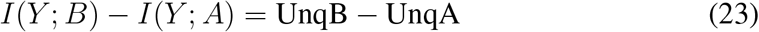

The term in (23) defines the unique information asymmetry. A positive value suggests that the information processing is being driven mainly by input *B*, whereas a negative value suggests that input *A* provides more drive.

When the partial information components are known *a priori* to be non-negative, we may deduce the following from (11), (18), (19). When the interaction information in (16) is positive, a lower bound on the synergy of a system is given by the interaction information (McGill, 1954). Also, the expression in (23) provides a lower bound for UnqB, when *I*(*Y* ; *B*) > *I*(*Y* ; *A*). Thus some deductions can be made without considering a PID. While such deductions can be useful in providing information bounds, it is only by computing a PID that the actual values of the partial information components can be obtained.

When making comparisons between different systems it is sometimes necessary to normalise the information decomposition by dividing each term by their sum, the output entropy, *H*(*Y*). Such normalisation will be applied in the analyses when comparing decompositions obtained under different conditions of an experimental or grouping factor. For probability distributions in which the inputs *B* and *A* are marginally independent the source shared information, ShdS, should be equal to zero, and hence the total shared information is entirely mechanistic shared information - shared information due to the probabilisitic mechanism involved in the information processing.

### A2. Derivation of transfer functions for burst firing

A transfer function for the probability of burst firing, *P*_2_(*b, a*), taking the strength (amplitude) of basal (somatic), *b*, and apical, *a*, current injections as arguments, was arrived at via the following reasoning and inspection of burst probability data obtained from simulations. To facilitate unpicking functional components contributing to bursting output we took a staged approach of examining a variety of bursting data, firstly from the HL regime in which apical input alone did not cause bursting. Here, varying the duration of the basal input helped reveal distinct contributions of basal and apical input. The transfer function derived from this data was then readily extended to cover the full HH range of input amplitudes.

#### Transfer function for HL regime with different basal stimulus durations

The initial data set of simulated spiking output covers an apical input range from 0 to 1 nA, and basal ranges dependent on the duration of the basal stimulus, being 0 to 3 nA for durations of 2 or 5 ms, and 0 to 1 nA for 10 ms duration. Apical input alone very rarely produces spiking output in this range. The basal input alone can produce spiking and for the 5 and 10 ms durations can also produce bursting at the higher end of their ranges. Burst probabilities were calculated from the frequency of output bursts (2 or more spikes with interspike intervals less than 25 ms) over 100 simulations for each combination of amplitudes of basal and apical input for discrete increments in their respective amplitudes within their defined ranges (see Section 2). This yielded a matrix of 31 by 11 probabilities for basal durations of 2 or 5 ms, and 21 by 11 for a basal duration of 10 ms. Contour maps of the burst probability across these different combinations of basal and apical input amplitudes and selected durations of basal input are shown in Figure 8. Plots of burst probability as a function of the strength of basal input, for different selected values of the apical input and durations of basal input, are shown in Figure 9 (dotted lines).

**Figure 8.**
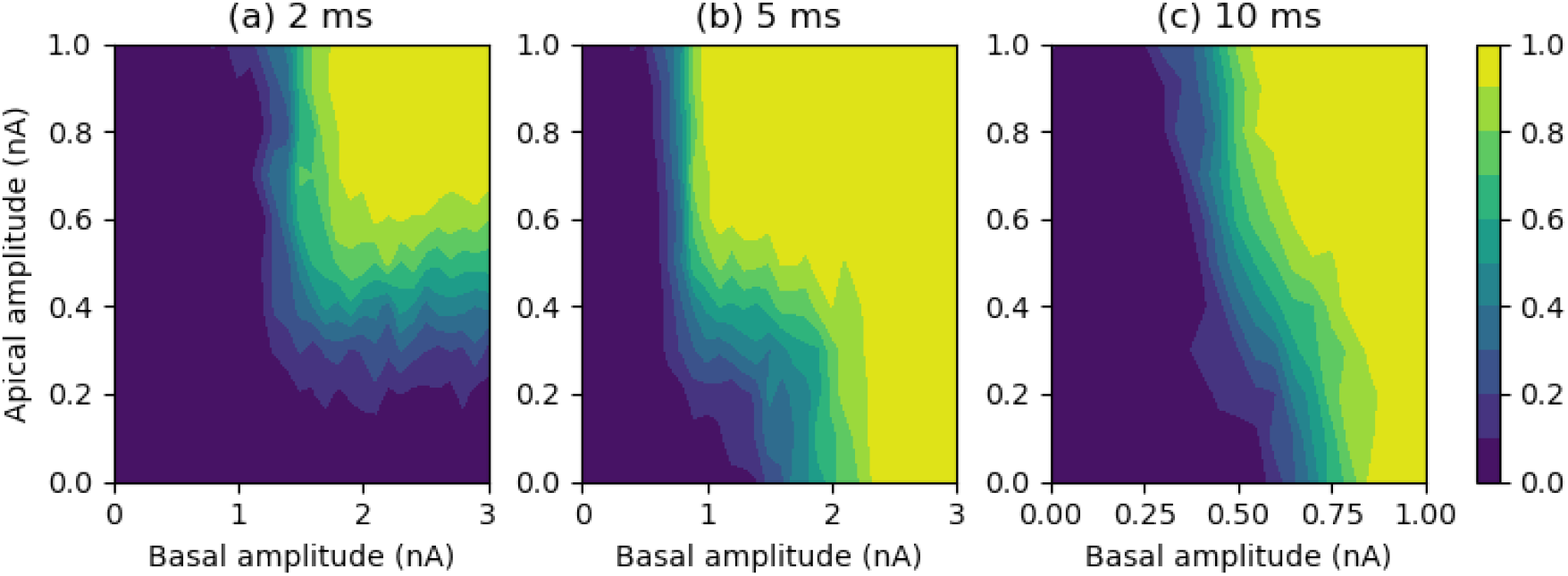
Contour maps of burst probability for different durations of basal input. Left: basal duration 2 ms, basal maximum amplitude 3 nA; Middle: 5 ms, max amp 3 nA; Right: 10 ms, max amp 1 nA; apical maximum amplitude is 1 nA in all cases.

**Figure 9.**
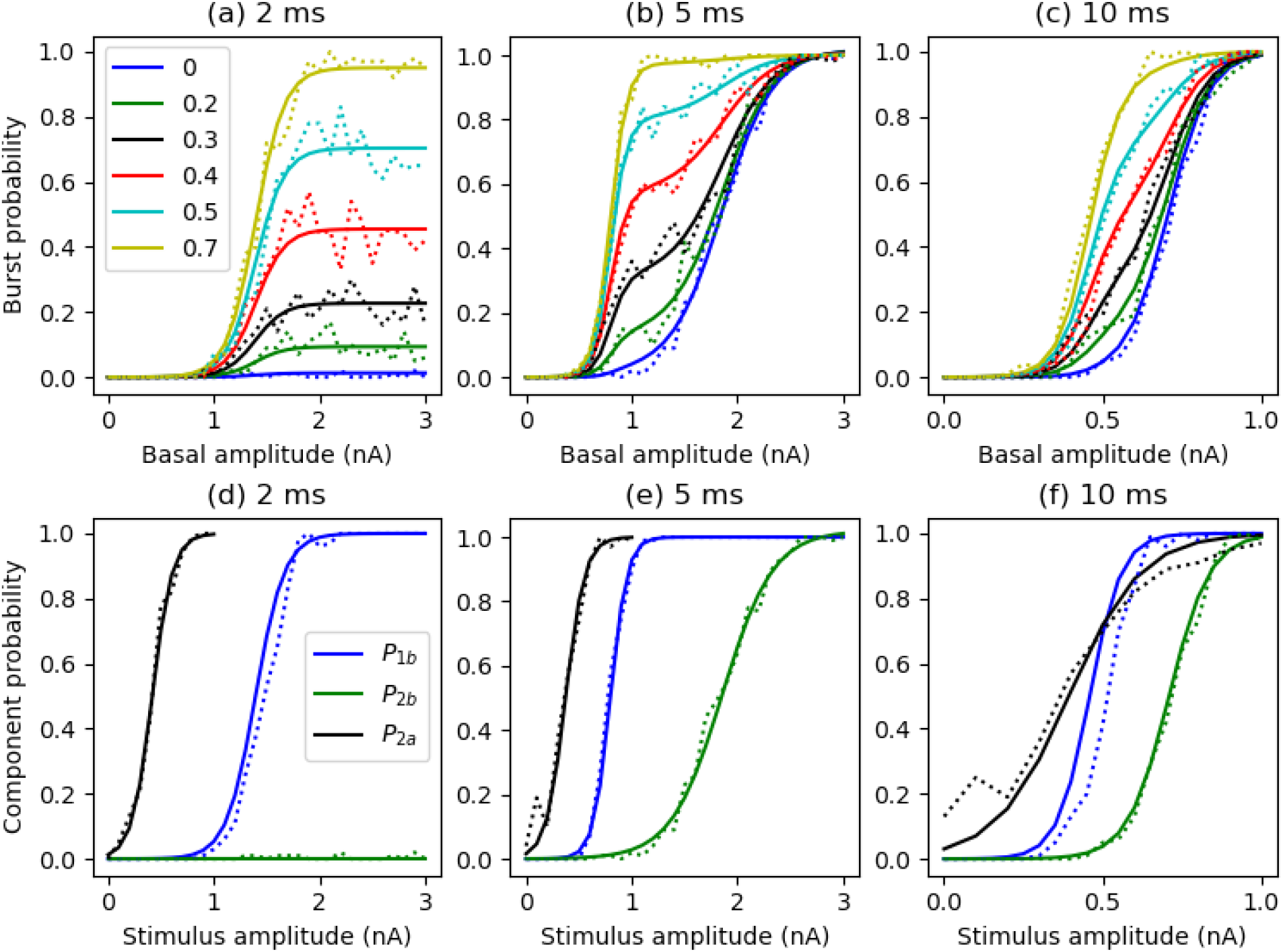
(a-c) Least-squares fit of transfer functions (solid lines), shown as burst probability versus basal amplitude, with plots for selected values of the apical amplitude (legend); dashed lines are the frequency data from simulations. (d-f) Comparison of transfer function components with estimates from the simulation data; first spike probability (*P*_1*b*_) and burst probability when apical amplitude is zero (*P*_2*b*_) are extracted exactly from the data; burst probability due to apical input alone (*P*_2*a*_) is estimated by data slices for particular basal amplitudes (b=2 nA for 2 ms duration, 1.2 nA for 5 ms and 0.6 nA for 10 ms). Left: basal duration 2ms; Middle: 5ms; Right: 10ms.

From the maps it can be seen that for longer duration basal inputs (5 or 10 ms) burst probability is an increasing function of basal amplitude, with high burst probability being achieved at lower basal amplitudes as the amplitude of the apical input is increased. For brief 2 ms basal inputs, once the basal input is sufficiently strong so as to produce a somatic spike then burst probability becomes an increasing function of the apical amplitude. Apical input alone produces a very low (essentially zero) burst probability over the range of amplitudes used.

Firstly, the range of apical input used cannot cause spiking output on its own, so the probability of burst firing factors as the product of the probability of a first spike, which is a function, *P*_1*b*_(*b*), of basal input only, multiplied by the probability of a second or more spikes given that a first spike has occurred, *P*_21_(*b, a*), which is a function of both basal and apical input. So we have:

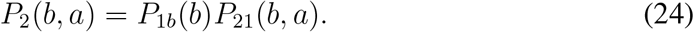

First spike probability is reasonably a sigmoidal function of basal amplitude as this basal input sums with the noisy (random) membrane current of mean zero, giving:

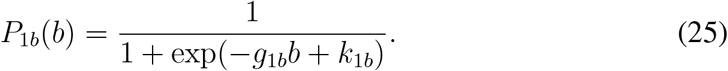

Examining the burst probability curves when basal duration is short (Figure 9a), shows that in this case, once a first spike has occurred then the probability of a burst is constant with increasing basal amplitude, but with this constant increasing with apical amplitude. This gives:

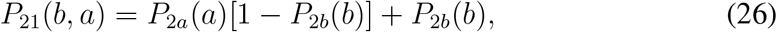

where *P*_2*b*_(*b*) is the burst probability achieved when apical input is zero; and *P*_2*a*_(*a*) is the probability of a burst due to apical input following a first somatic spike. The burst probabilities reached with increasing *a* are well matched by *P*_2*a*_(*a*) being a sigmoidal function of *a*:

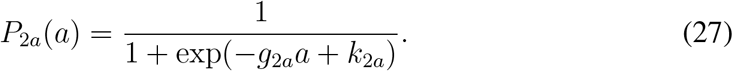

Finally, when basal duration is longer than 2ms, basal input alone is able to generate bursts with increasing probability with increasing basal amplitude. This increasing burst probability clearly has a sigmoidal relationship with basal amplitude (Figure 9b,c blue lines), so we define:

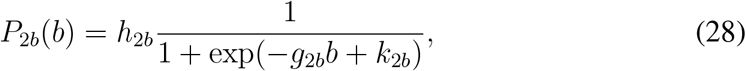

where scaling factor *h*_2*b*_ ⪡ 1 when basal duration is 2ms, but *h*_2*b*_ ≈ 1 for longer basal durations.

The complete HL transfer function is thus given by the following set of equations:

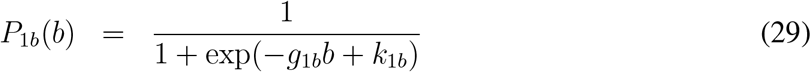

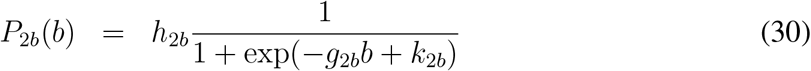

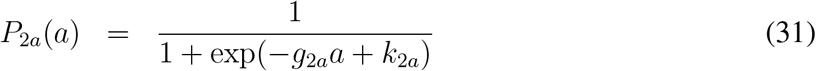

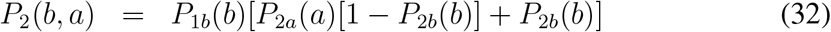

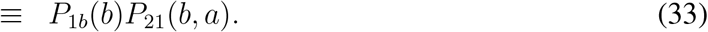

The least-squares fits of this transfer function, *P*_2_(*b, a*), to the simulation data for different basal durations are illustrated in Figure 9a-c. The corresponding transfer function parameter values are given in Table 3.

**Table 3:** Least squares fits of *P*_2_(*b, a*) transfer function parameter values for burst firing with different durations of basal input (2, 5 and 10 ms). Values are plus/minus their estimated standard error (apart from the 2*b* parameters at 2 ms duration, which are not well constrained by the data, but where the scaling factor *h*_2*b*_ is very small).

| Duration: | 2 ms | 5 ms | 10 ms |
| --- | --- | --- | --- |
| $h_{2b}$ | 0.0019 | $1.02 \pm 0.01$ | $1.0 \pm 0.014$ |
| $g_{2b}$ | 0.0012 | $4.15 \pm 0.16$ | $15.43 \pm 0.82$ |
| $k_{2b}$ | 40.4 | $7.7 \pm 0.29$ | $10.94 \pm 0.57$ |
| $g_{1b}$ | $7.3 \pm 0.41$ | $12.66 \pm 0.69$ | $19.81 \pm 1.14$ |
| $k_{1b}$ | $10.18 \pm 0.57$ | $10.09 \pm 0.54$ | $9.09 \pm 0.5$ |
| $g_{2a}$ | $10.45 \pm 0.28$ | $11.06 \pm 0.37$ | $8.8 \pm 0.49$ |
| $k_{2a}$ | $4.36 \pm 0.12$ | $4.13 \pm 0.15$ | $3.46 \pm 0.19$ |

To support the veracity of this transfer function, the correspondence between the different components and aspects of first spike and bursting probability data is shown in Figure 9d-f. Here, *P*_2*b*_ is a direct fit to the underlying data, whereas *P*_1*b*_ and *P*_2*a*_ emerge implicitly from the optimisation and are not explicit fits, but nonetheless show a strong correspondence to the equivalent data. Note that the data shown for the first spike probability, *P*_1*b*_, has been extracted from the simulation data, but is not explicitly used in the optimisation. The contribution of the apical amplitude to bursting, *P*_2*a*_, is not explicitly available in the data, but good estimates are available using slices through the data for basal amplitude values that give a very low burst probability when the apical amplitude is zero, but have a burst probability near one when combined with strong apical input; these are the data examples shown in Figure 9d-f.

#### Full HH regime transfer function

The transfer function above (Equation 32), that covers the two regimes with low apical input (LL and HL), can be extended to cover the other two regimes (LH and HH) in which strong apical inputs alone can generate output bursting by including a factor that accounts for such bursting. The resultant transfer function is the original function fractionally summed with the contribution to bursting by apical input alone:

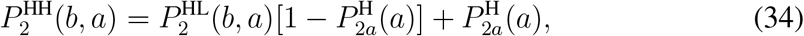

where 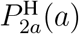 is a sigmoidal function of the apical amplitude (with superscript H designating the high apical range only):

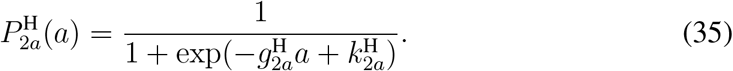

This can be rearranged to yield a form that is a sum of terms that depend on the basal and apical inputs alone plus a contribution due to the apical and basal inputs combined:

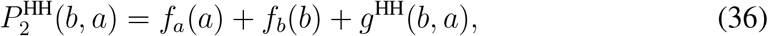

where 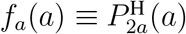 and 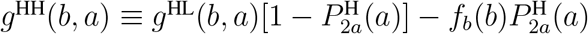.

We fit this transfer function, 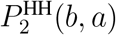, to the full data set for a basal duration of 10 ms. A least squares fit of 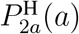 to the collected burst probabilities when the basal input is 0 yields 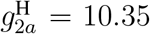 and 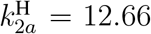. Using these parameter values, the full HH transfer function fits the data well across the entire range of basal and apical input amplitudes (regime HH), as illustrated in Figure 10. This reveals that the influence of basal on apical input and vice versa is now qualitatively symmetric, with both being able to modulate the effect of the other (see also Figure 4d). However, the quantitative modulation is different, as indicated by the shape of the curves in Figure 10a compared to Figure 10b. In neither case is the modulation simply a change in gain or bursting threshold (shift), rather the ‘modulating’ input increases the likelihood of bursting over a range of the ‘driving’ input.

**Figure 10.**
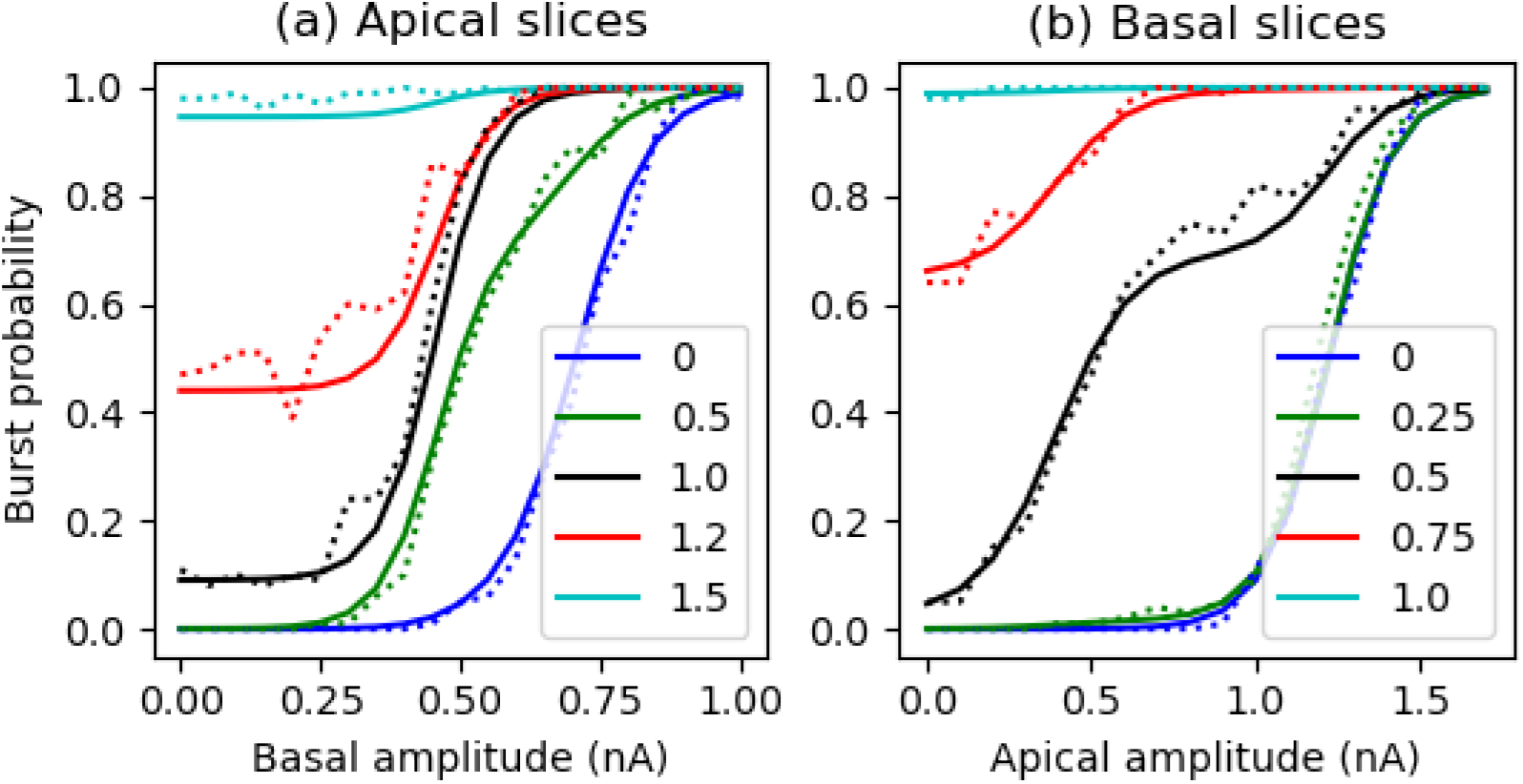
Fit of full transfer function to HH data set for a basal duration of 10 ms. (a) Transfer function for selected values of the apical amplitude (solid lines) plotted against the simulated burst probabilities (dotted lines) across the range of basal amplitudes. (b) Transfer function for selected values of the basal amplitude (solid lines) plotted against the simulated burst probabilities (dotted lines) across the range of apical amplitudes.

In the regime of low basal but high apical input, in which basal input alone does not produce a burst (*f*_*b*_(*b*) ≈ 0), this full transfer function collapses to 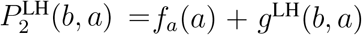, where 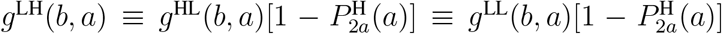 where *g*^HL^(*b, a*) reduces to *g*^LL^(*b, a*) ≡ *P*_1*b*_(*b*)*P*_2*a*_(*a*) since *P*_2*b*_(*b*) ≈ 0 in this regime.

